# AIM2 inflammasome activation in astrocytes occurs during the late phase of EAE

**DOI:** 10.1101/2021.10.03.462457

**Authors:** William E. Barclay, M. Elizabeth Deerhake, Makoto Inoue, Toshiaki Nonaka, Kengo Nozaki, Nathan A. Luzum, Nupur Aggarwal, Edward A. Miao, Mari L. Shinohara

**Author notes:** **Corresponding Author:** Mari L. Shinohara.

## Abstract

Inflammasomes are a class of innate immune signaling platforms that activate in response to an array of cellular damage and pathogens. Inflammasomes promote inflammation under many circumstances to enhance immunity against pathogens and inflammatory responses through their effector cytokines, IL-1β and IL-18. Multiple sclerosis and its animal model, experimental autoimmune encephalomyelitis (EAE), are such autoimmune conditions influenced by inflammasomes. Despite work investigating inflammasomes during EAE, little remains known concerning the role of inflammasomes in the central nervous system (CNS) during the disease. Here we use multiple genetically modified mouse models to monitor activated inflammasomes *in situ* based on ASC oligomerization in the spinal cord. Using inflammasome reporter mice, we found heightened inflammasome activation in astrocytes after the disease peak. In contrast, microglia and CNS-infiltrated myeloid cells had few activated inflammasomes in the CNS during EAE. Astrocyte inflammasome activation was dependent on AIM2, but low IL-1β expression and no significant signs of cell death were found in astrocytes during EAE. Thus, the AIM2 inflammasome activation in astrocytes may have a distinct role from traditional inflammasome-mediated inflammation.

**SIGNIFICANCE STATEMENT:** Inflammasome activation in the peripheral immune system is pathogenic in multiple sclerosis (MS) and its animal model, experimental autoimmune encephalomyelitis (EAE). However, inflammasome activity in the central nervous system (CNS) is largely unexplored. Here, we used genetically modified mice to determine inflammasome activation in the CNS during EAE. Our data indicated heightened AIM2 inflammasome activation in astrocytes after the disease peak. Unexpectedly, neither CNS-infiltrated myeloid cells nor microglia were the primary cells with activated inflammasomes in SC during EAE. Despite AIM2 inflammasome activation, astrocytes did not undergo apparent cell death and produced little of the proinflammatory cytokine, IL-1β, during EAE. This study showed that CNS inflammasome activation occurs during EAE without associating with IL-1β-mediated inflammation.

## INTRODUCTION

Multiple sclerosis (MS) and its mouse model, experimental autoimmune encephalomyelitis (EAE), are demyelinating neurodegenerative diseases punctuated by inflammatory immune reactions in the central nervous system (CNS). Inflammasomes are sensors of a wide range of microbe-associated molecular patterns (MAMPs) and damage-associated molecular patterns (DAMPs) and induce inflammation (1). In EAE, the NLRP3 inflammasome has a particularly well-documented role in the peripheral immune system in promoting immune cell recruitment to the CNS through the generation of the inflammatory cytokines IL-1β and IL-18 by peripheral myeloid cells (2-5). However, despite much work centered on inflammasomes in the peripheral immune response, the role of inflammasomes in the CNS is significantly less understood.

Inflammasomes are distinct from other PRRs in mode of signaling and downstream effector function. First, a sensor (*e*.*g*., NLRP3, AIM2) forms a scaffold, to which the inflammasome adaptor ASC binds and polymerizes (called the “ASC speck”). Pro-caspase-1 associates with the polymer, then self-cleaves to become proteolytically active caspase-1, which further activates downstream substrates, including pro-IL-1β, pro-IL-18, and the pore-forming protein, gasdermin-D (GSDMD). Cleaved GSDMD forms a pore in cellular membranes and induces pyroptosis, resulting in a release of mature IL-1β and IL-18 to the extracellular space. In MS, genetic variation in inflammasome signaling pathways was reported (6, 7). Particularly, the NLRP3 inflammasome was demonstrated to be a prognostic factor and a therapeutic target in primary progressive MS (8). In EAE, NLRP3 and ASC are necessary for passive and standard (“Type-A”) active EAE (2-4). NLRP3 inflammasome activation in macrophages and dendritic cells in secondary lymphoid organs results in IL-1β and IL-18 release, which induces expression of chemokines and their receptors required for leukocyte CNS entry (2). We have also demonstrated that the NLRP3 inflammasome can be dispensable for EAE induction if the innate immune system is strongly activated with aggressive immunization schemes (“Type-B EAE”)(9).

As inflammasome activation is a post-translational process, the expression of inflammasome components does not necessarily indicate their activation. Thus, separately assessing both expression and activation of inflammasomes is critical. Recent studies indicated that microglia, astrocytes, and neurons can activate inflammasomes; and most identification was performed *ex vivo* (8, 10-17). However, defining inflammasome activation *in situ* is critical because CNS-resident cells significantly alter their behavior once isolated from tissues. So far, only a limited number of studies has demonstrated unequivocal activation of inflammasomes *in situ* in the CNS (10-12).

In this report, we identified activated inflammasomes in the CNS during EAE using the ASC-Citrine mouse line, which allows *in situ* detection of activated inflammasomes (18). Our study identified maximal inflammasome activation in the spinal cord (SC) after the EAE peak, which contrasts with the inguinal lymph nodes (iLNs), in which inflammasome activation was present at pre-symptomatic disease. Unexpectedly, neither microglia nor CNS-infiltrated myeloid cells were the primary cells with activated inflammasomes in SC during EAE. Instead, we detected inflammasome activation mainly in astrocytes and limited inflammasome activation in motor neurons. Furthermore, we found that the AIM2 inflammasome is activated in astrocytes during EAE. However, even with AIM2 inflammasome activation, astrocytes did not clearly undergo cell death and have poor *Il1b* gene expression, suggesting the possibility that AIM2 inflammasome activation in astrocytes serves a different purpose than traditional inflammasome-mediated inflammation.

## RESULTS

### Inflammasome activation in the spinal cord at a late stage of EAE

Inflammasome signaling is critical to EAE development in the peripheral lymphoid organs (2-5, 9). Yet, the extent and spatiotemporal distribution of inflammasome activation in the CNS during EAE is largely unknown. Classically, detection of inflammasome activation is performed by identifying cleaved caspase-1 by Western blotting (WB), which cannot be applied *in situ*. Therefore, we used a different molecular signature of inflammasome activation – the oligomerization of ASC, microscopically observed as the “ASC speck.” In this study, we used inflammasome activation reporter mice, which express ASC fused to a fluorescent Citrine protein (ASC-Citrine)(18). The ASC-Citrine reporter allows *in situ* detection of active inflammasomes by visualization of ASC specks (12, 18) (**Fig. S1*A***), and the presence of the reporter did not alter disease course of EAE (**Fig. 1*A***). The ASC-Citrine reporter was validated for use in tissue by immunostaining against ASC in live spleen slice cultures following NLRP3 inflammasome activation (**Fig. S1*A***).

**Figure 1.**
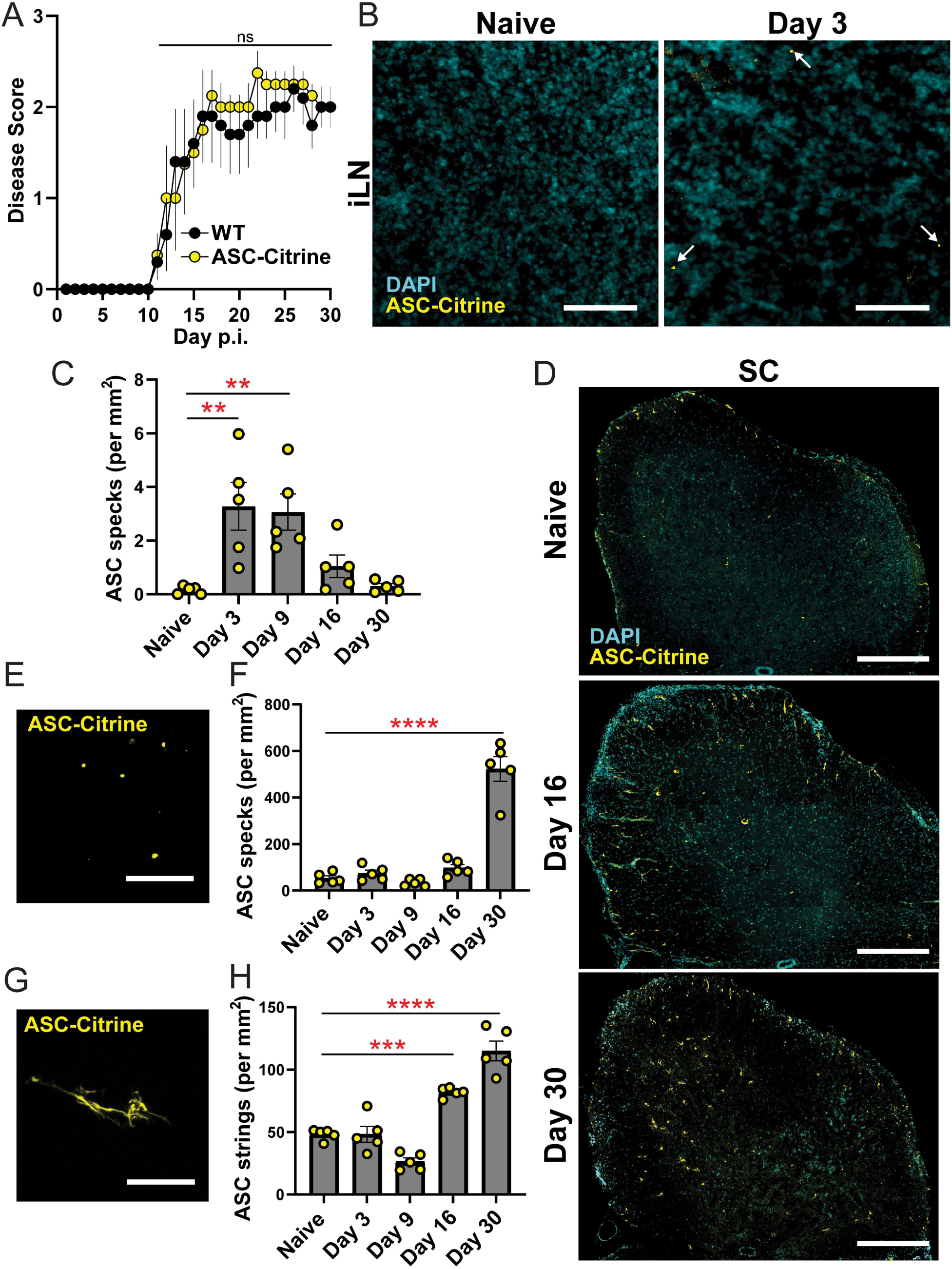
Inflammasome activation in the CNS during late EAE. **(A)** EAE disease score of WT (*n*=5) vs. ASC-Citrine mice (*n*=4). Mann-Whitney test of total AUC for disease was used. **(B, C)** Representative images *(B)* and quantification *(C)* of ASC specks in the iLNs of ASC-Citrine mice during EAE. Each datapoint represents a value of an average value from two cross-sections of both iLNs (25 μm thickness) per mouse. *n=5* mice. One-way ANOVA, *p*=0.0006, with Dunnett’s multiple comparisons test. Scale bar is 20 μm. **(D)** Representative images of SC from ASC-Citrine mice at indicated time points during EAE. Scale bar is 300 μm. **(E-H)** Representative image *(E, G)* and quantification *(F, H)* of ASC specks *(E, F)* and ASC strings *(G, H)* in SC from ASC-Citrine mice during EAE. Scale bar is 10 μm. Each datapoint represents a value from one mouse (*n*=5). Two coronal cross-sections (25 μm thickness) of L5 SC from one mouse were quantified manually and averaged. One-way ANOVA with Dunnett’s multiple comparisons tests *(C, F, H)*. ^ns^ *p*>0.05, \*\**p* < 0.01, \*\*\**p*<0.001, \*\*\*\**p*<0.0001. Error bars denote mean ± SEM.

Before evaluating inflammasome activation in the CNS, we first visualized and quantified ASC specks in the iLNs as the primary site of immune reaction to EAE induction. ASC specks were detected at 3 days post induction (dpi) of EAE (**Fig. 1*B, C***), which is well before the disease onset, and continually detected until 9-dpi (**Fig. 1*C***). However, ASC specks were almost undetectable by the point of disease peak (16-dpi) and after (**Fig. *1C***). The cervical lymph nodes (cLN) have also been noted as a site of primary immune reaction in some models of EAE (19, 20), but few ASC specks were detected there throughout disease (**Fig, S1B**). The spinal cord (SC) of ASC-Citrine mice exhibited a much higher number of ASC specks than the iLNs (**Fig. 1*D*-*F***) with a significant increase in the number of ASC specks in the later phase of EAE at 30-dpi (**Fig. 1*D, F***). Further, in addition to ASC specks, we also observed atypical fiber-like ASC-Citrine signals, which we termed “ASC strings,” unique to the CNS (**Fig. 1*G***). While different in magnitude, both ASC specks and ASC strings appeared with the largest increases at the later phase of disease, around 30-dpi (**Fig. 1*F, H***). Due to the abundance of ASC specks and strings at 30-dpi, all subsequent analyses in SC were at 30-dpi, unless otherwise stated. This quantification was also performed manually, and all subsequent ASC speck and string quantification was performed using the Imaris software.

We have previously shown a sub-type of EAE (Type-B EAE) which does not require the NLRP3 inflammasome in the peripheral lymphoid organs to develop EAE (9). In the CNS, Type-A and Type-B EAE had comparable numbers of ASC specks (**Fig. S1*C, D***), suggesting the more consistent connection of EAE severity and active inflammasome in the CNS than in the periphery. In sum, these results suggest that inflammasomes are activated in the SC during both Type-A and Type-B EAE and that their activation is heightened after peak disease.

### Inflammasome activation in non-BM-derived cells in the spinal cord

Because we identified inflammasome activation in the CNS during EAE, the role of ASC in non-hematopoietic cells was investigated with a bone marrow (BM) chimera approach using CD45.1 congenic donor mice. We compared two groups of BM chimeras, generated by adoptively transferring WT BM cells to either WT or *Pycard*^-/-^ (ASC knockout) recipients (the extent of reconstitution is shown in **Fig. S2*A***). Compared to WT recipients, *Pycard*^*-/-*^ recipients demonstrated milder EAE after the disease peak (**Fig. *2A-C***), suggesting that ASC in non-hematopoietic cells impacted EAE severity after the disease peak. Next, we sought to determine if CNS-infiltrated cells possess activated inflammasomes in the SC during EAE. Two groups of BM chimeras were compared; one group with ASC-Citrine BM donor cells to wild-type (WT) recipients and the other with WT BM cells to ASC-Citrine recipients. Reconstitution of approximately 90% of BM cells were confirmed (**Fig. S2*B***) and no impact of ASC-Citrine expression on EAE development was confirmed (**Fig. S2*C***). ASC specks were identified in the iLNs of WT recipients reconstituted with ASC-Citrine BM (**Fig. 2*D, E***). However, unexpectedly, the mice showed no ASC specks and strings in the SC (**Fig. 2*F-H***; **S2*D***). In contrast, ASC specks and strings were identified in the SC of ASC-Citrine recipients transferred with WT BM cells (**Fig. 2*F-H***). This suggests that the source of inflammasome activation is CNS-resident cells in the SC.

**Figure 2.**
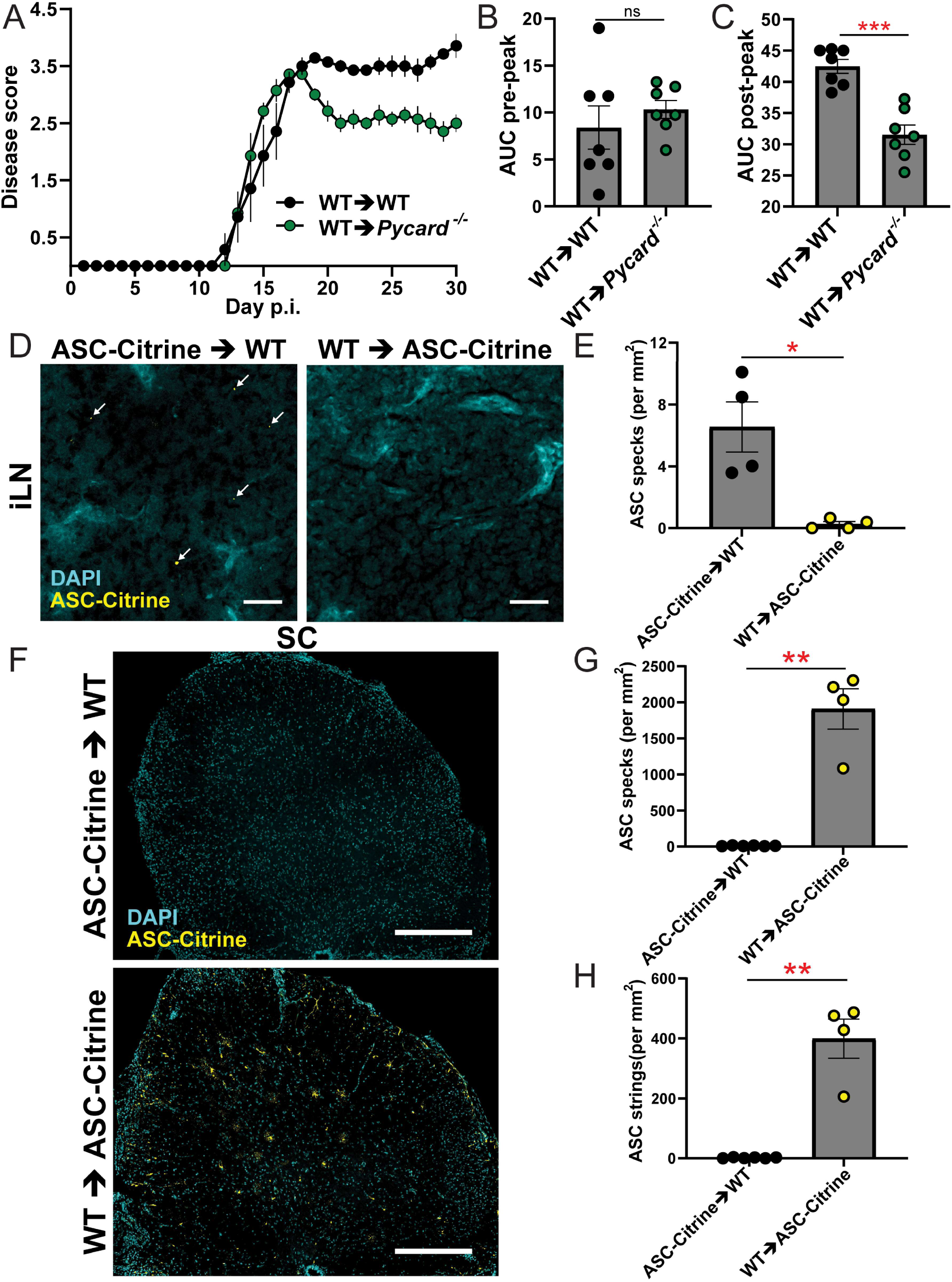
No inflammasome activation in hematopoietic cells in the CNS during EAE. **(A)** EAE disease scores of WT (BM donor)→WT (recipient) chimeras vs. WT→*Pycard*^*-/-*^ chimeras. *n*=7, combined from multiple experiments. **(B, C)** Comparison of EAE disease severity of WT→WT chimeras versus WT→*Pycard*^*-/-*^ chimeras. Each datapoint represents a value from one mouse (*n=*7), combined from multiple experiments. Area under curve (AUC) quantification of pre-peak disease *(B)*, AUC quantification of post-peak disease *(C)*. **(D, E)** Representative images *(D)* and quantification *(E)* of ASC specks in the iLNs of WT→ASC-Citrine chimeras versus ASC-Citrine→WT chimeras at 3 dpi of EAE. Each datapoint represents a value from one mouse as an average of both iLNs (*n*=4). Mann-Whitney test was used. Scale bar is 20 μm. **(F-H)** Representative images *(F)* and quantification of ASC specks *(G)* and ASC strings *(H)* of SC from WT→ASC-Citrine BM chimeras (*n*=4) versus ASC-Citrine→WT chimeras (*n*=6) at 30-dpi of EAE. Each datapoint represents a value from one mouse. Mann-Whitney test was used (*B, C, E, G, H*). Scale bar is 300 μm. *(B, C, G, H)*. ^ns^ *p*>0.05, \**p*<0.05, \*\**p*<0.01, \*\*\**p*<0.001. Error bars denote mean ± SEM.

### Inflammasome activation in CNS during EAE in astrocytes

We next sought to identify CNS-resident cells with activated inflammasomes during EAE. ASC specks and strings were identified, and then cell types were assigned by counterstaining to identify microglia (TMEM119), astrocytes (GFAP or ALDH1L1), neurons (NeuN), oligodendrocyte precursor cells (NG2), or mature oligodendrocytes (MBP) (**Fig. 3*A-C*; S2*E, F***). The majority of ASC specks and strings were found in astrocytes, while a small number were classified as microglial and neuronal (**Fig. 3*D*-*F***). Few ASC specks or strings were detected in oligodendrocyte precursor cells (OPCs) or mature oligodendrocytes (**Fig. S2*G, H***). In neurons, all ASC specks were found in cell bodies of ChAT^+^ alpha motor neurons (ChAT^+^NeuN^+^) in the ventral horn (VH), but these neuronal ASC specks did not increase during EAE (**Fig. S2*I***).

**Figure 3.**
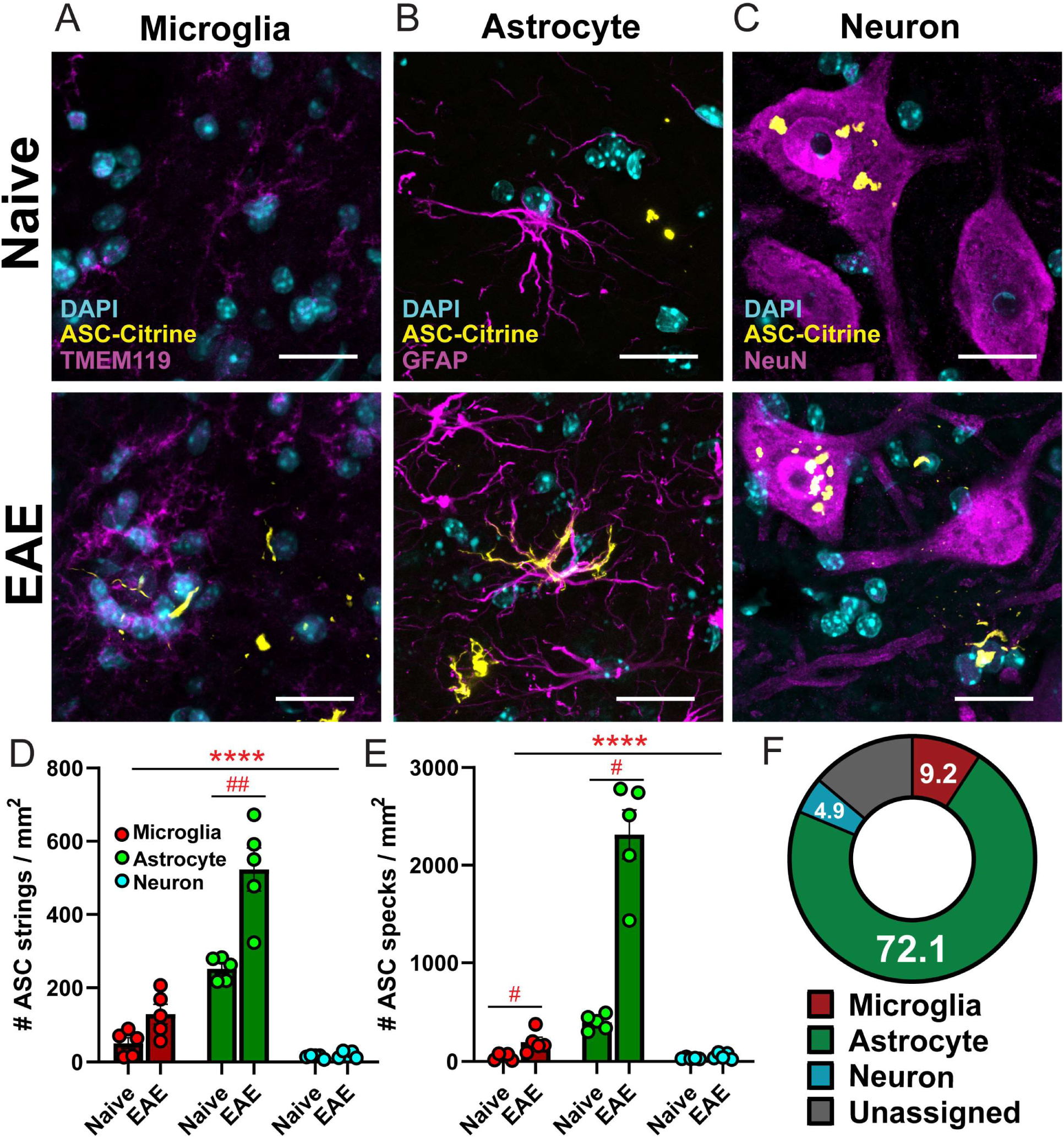
Identifying inflammasome activation in CNS cells with an immunofluorescent approach. **(A-C)** Representative images of ASC specks and strings counter-stained with antibodies against TMEM119 for microglia *(A)*, GFAP for astrocytes *(B)*, and NeuN for neurons *(C)* in SC from naïve versus 30-dpi EAE ASC-Citrine mice. Scale bar is 20 μm. **(D, E)** Quantification of ASC specks *(D)* and ASC strings *(E)* in microglia, astrocytes, and neurons in the ventral horn (VH) of SC from naïve versus 30-dpi EAE ASC-Citrine mice. Two-way repeated measures (RM) ANOVAs were used (main effect of cell type: \*\*\*\**p*<0.0001 (D, E)), with Sidak’s multiple comparisons test post hoc (^#^*p*<0.05, ^*##*^*p*<0.01). **(F)** Relative contribution of microglia, astrocytes and neurons to total number of ASC specks in L5 SC at 30-dpi EAE. “Unassigned” indicates unindentified cell sources of ASC-Citrine signals. Each datapoint represents a value from one mouse (*n*=5). Two-way RM ANOVA was used (main effect of cell type: *p*<0.001) with Sidak’s multiple comparisons test post hoc. \*\**p*<0.01. Error bars denote mean ± SEM.

We further validated these findings with a cell type-specific ASC-Citrine reporter approach by using *Asc*-*Citrine*^*LSL*^ mice, which retain an LSL cassette upstream of the ASC-Citrine construct. To express ASC-Citrine in astrocytes and neurons in a cell type-specific manner, we used *Gfap*^*Cre*^*;Asc*-*Citrine*^*LSL*^ and *Syn1*^*Cre*^*;Asc*-*Citrine*^*LSL*^ mice, respectively. For microglia-specific ASC-Citrine expression, we used *Cx3cr1*^*CreERT2*^*;Asc*-*Citrine*^*LSL*^ mice treated with tamoxifen (TAM) with a six-week “wash out” period to exclude ASC-citrine expression in myeloid cells other than microglia (**Fig. S3*A***) by taking an advantage of the long half-life of microglia (gating strategy to evaluate microglia is shown in **Fig. S3*B***). We considered that using the microglia-specific ASC-Citrine reporter mice was critical because the expression level of Tmem119, used in **Fig. 3*A-F***, decreases as EAE progresses (21) and may confound some image analysis. No alteration in EAE severity was confirmed in the group of mutant mice expressing Cre in targeted cell types (**Fig. S3*C-E***). The SC of these mutant mice was analyzed during EAE by confocal microscopy (**Fig. 4*A-C***). Again, a high number of ASC specks and strings were confirmed in astrocytes by using the *Gfap*^*Cre*^*;Asc*-*Citrine*^*LSL*^ mice, while few ASC specks or strings were identified in microglia (**Fig. 4*D, E***). Neurons also showed a consistent but small number of ASC specks (**Fig. 4*D, E***). Nonetheless, these data mirrored the results by antibody counterstaining in **Fig. 3*A-G***, strongly indicating that inflammasome activation is predominantly in astrocytes during EAE.

**Figure 4.**
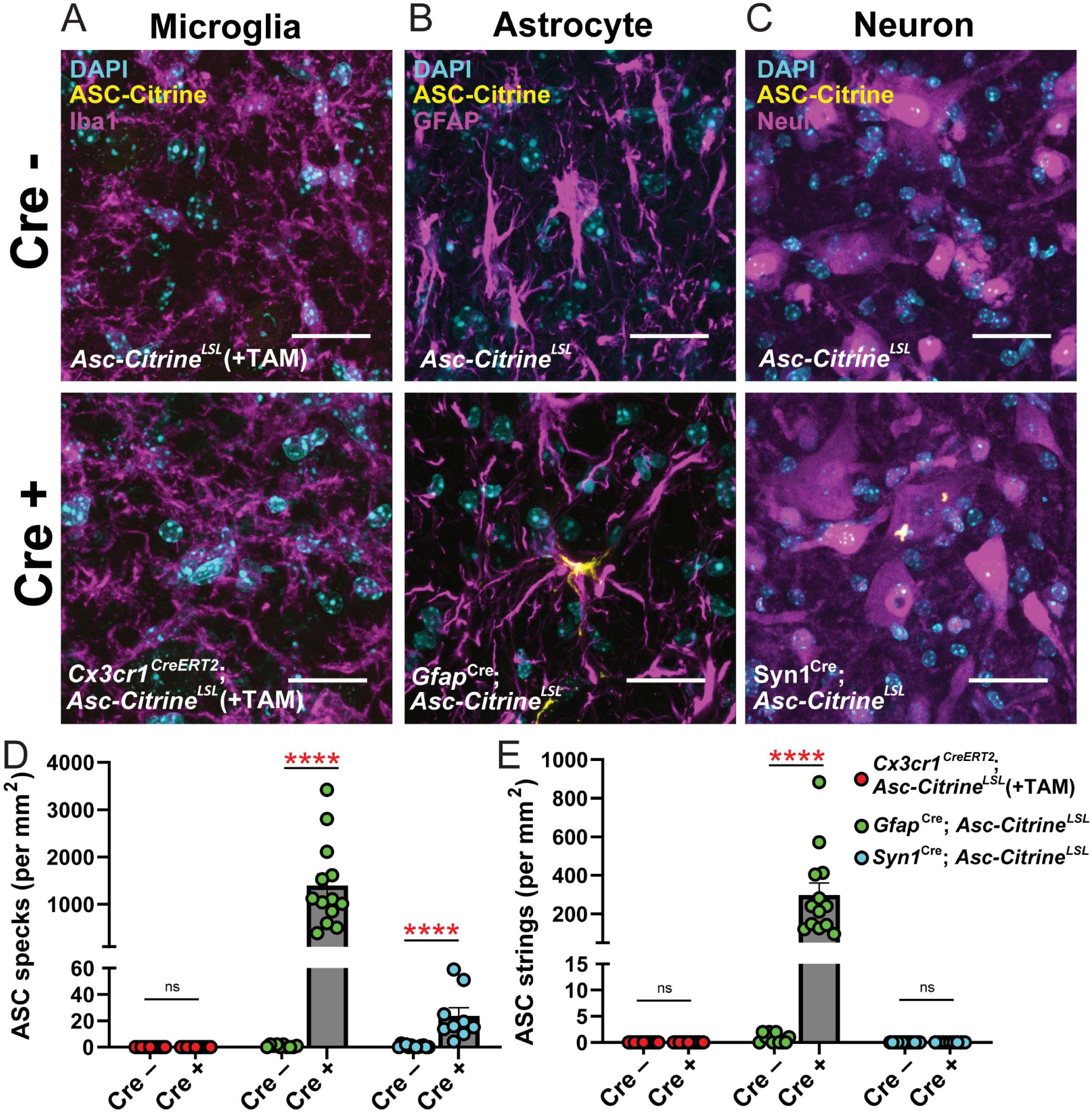
Identifying Inflammasome activation in CNS cells by a mouse genetics approach. **(A-C)** Representative images of SC in *Asc-Citrine*^*LSL*^ versus *Cx3cr1*^*CreERT2*^;*Asc-Citrine*^*LSL*^ mice *(A), Gfap*^*Cre/+*^;*Asc-Citrine*^*LSL*^ mice *(B)*, and *Syn1*^*Cre/+*^*;Asc-Citrine*^*LSL*^ mice *(C)* at day 30-dpi of EAE. Mice for microglia evaluation were treated with tamoxifen. Scale bar is 20 μm. **(D, E)** Quantification of ASC specks *(D)* and ASC strings *(E)* in SC VH of mice indicated in (A-C). Each datapoint represents a value from one mouse. Combined from multiple experiments. *(D, E)* Mann-Whitney test was used. \*\*\*\**p*<0.0001. Error bars denote mean ± SEM.

### Limited induction of IL-1β-mediated inflammation by astrocytes during EAE

In EAE, astrocytes become activated in a process called astrogliosis; these activated cells gain a pro-inflammatory phenotype and are termed “reactive astrocytes” (22, 23). Here we investigated whether astrogliosis correlates with inflammasome activation. Astrogliosis was detected as increased GFAP intensity at 30-dpi (**Fig. 5*A, B***), but GFAP intensity did not correlate on a per cell basis with the presence of ASC specks (**Fig. 5*C, D***). We similarly found no correlation between inflammasome activation and individual acquisition of the neurotoxic “A1” reactive astrocyte phenotype (24, 25), based on the A1-astrocyte marker C3d (26), although C3d expression was enhanced in astrocytes in aggregate during EAE (**Fig. 5*C, E***).

**Figure 5.**
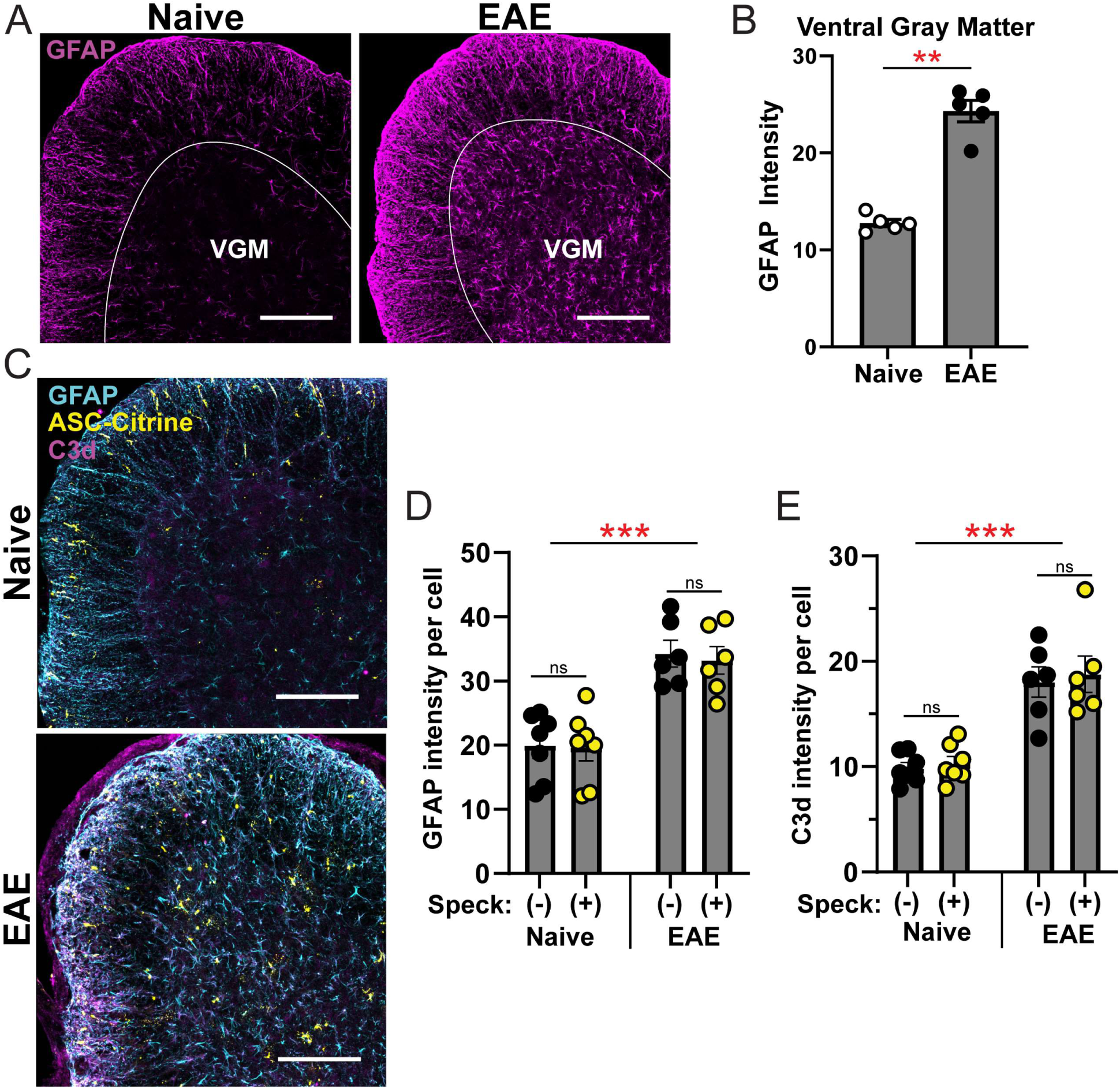
Evaluation of astrocytes with activated inflammasomes. **(A, B)** Representative images *(A)* and quantification *(B)* of total GFAP intensity in SC ventral gray matter from naïve 30-dpi EAE mice. Scale bar is 200 μm. GFAP intensity was quantified as mean signal intensity of GFAP in the ventral grey matter (VGM) using ImageJ. Mann-Whitney test was used. **(C-E)** Representative images *(C)* and quantification of GFAP intensity *(D)* and C3d intensity *(E)* per cell in gray matter SC astrocytes with and without ASC specks/strings from naïve ASC-Citrine (*n*=7) mice versus ASC-Citrine mice at 30-dpi of EAE (*n*=6). Scale bar is 200 μm. Each datapoint represents a value from one mouse. Individual astrocytes were identified using the Imaris software and the mean intensity per cell was quantified for GFAP and C3d. Two-way RM ANOVA was used with Sidak’s multiple comparisons test post-hoc. \*\**p*<0.01. \*\*\**p*<0.001. Error bars denote mean ± SEM.

To evaluate astrocyte gene expression during EAE, we first re-analyzed publicly available data obtained with a *Gfap*^*Cre*^-driven RiboTag mouse system, which allows purification of astrocyte-specific RNA (GSE100329)(27). Gene expression in total SC and SC astrocytes were compared between naïve and 30-dpi EAE mice. We found low expression of *Il1b* and genes encoding inflammasome sensor proteins (**Fig. S4*A***). Expression levels of *Il18* and *Casp1* were enriched in astrocytes independent of EAE (**Fig. S4*A***). To validate this data, we evaluated gene expression by RT-qPCR in total SC cells and astrocytes enriched by bead selection from naïve and 30-dpi EAE mice. Astrocyte-enriched cells showed generally low expression of genes encoding proteins related to inflammasomes even during EAE. Notably, the expression of *Il1b, Casp-1*, and *Gsdmd* was significantly lower in astrocyte-enriched cells than total SC cells (normalized to *Il1b* and *Casp1* expression in naïve total SC cells for **Fig. 6*A*** and ***B***, respectively). In the qPCR analyses, a majority of genes shown in **Fig. S4*A*** had mRNA levels that were close to the detection limit, despite reasonably high total RNA amounts, suggesting the general low expression of inflammasome-related genes in astrocytes. Under the low gene expression, we did not observe astrocyte enrichment of *Il18* and *Casp1* expression, as suggested in the RiboTag data (**Fig. S4*A*)**. This was consistent with the limited detection of the inflammasome-related proteins caspase-1, IL-1β, and GSDMD in SC astrocytes of either naïve mice or mice with EAE 30 dpi by immunostaining, despite robust detection in the spleen (**Fig. S4*B-G***). However, mild expression of GSDMD was detected in SC astrocytes following EAE induction (**Fig. S4D, G**). Next, we performed Western blotting (WB) to evaluate protein levels and inflammasome activation *in vitro*. Here, we used the C8-S astrocyte cell line due to the difficulty of culturing primary astrocytes, especially without altering their character in tissue culture settings. We compared C8-S to bone marrow-derived macrophages (BMDMs) as a positive control. We stimulated the NLRP3 or AIM2 inflammasome with nigericin or poly(dA:dT)-liposomes, respectively, after ultrapure LPS pre-treatment. Culture supernatants of C8-S showed a scarcity of cleaved caspase-1, IL-1β and IL-18 (**Fig. 6*C*; S5*A-C***). Notably, C8-S cell lysates also showed greatly reduced pro-caspase-1, pro-IL-1β, GSDMD-FL, and GSDMD-NT, compared to BMDMs (**Fig. 6*C*; S5*D-H***). These results suggest that astrocytes do not induce inflammation, mediated particularly by IL-1β as macrophages do.

**Figure 6.**
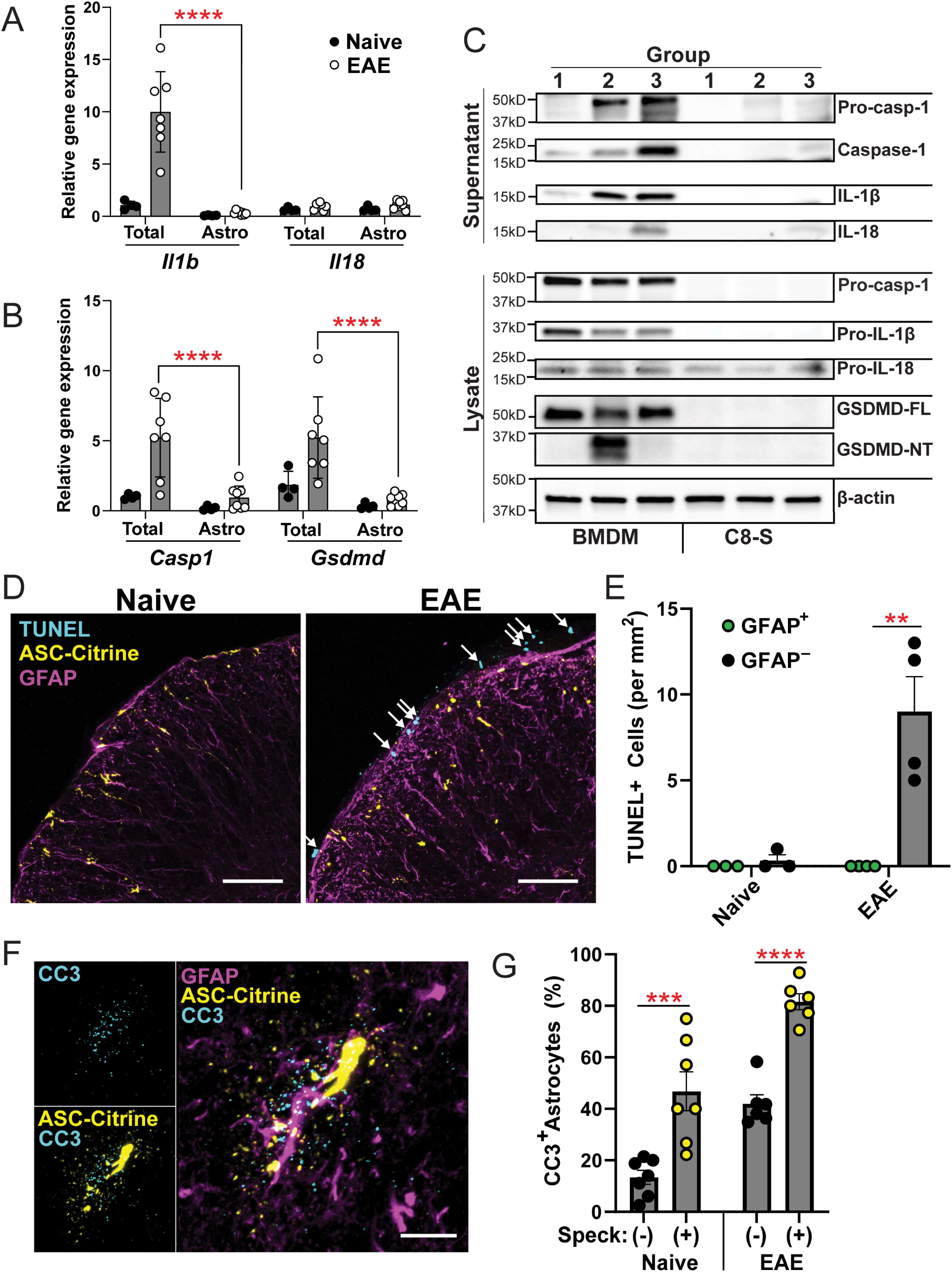
Outcomes of inflammasome activation in astrocytes. **(A, B)** RT-qPCR evaluation of inflammasome-related genes in SC total cells vs. astrocytes isolated from mice with EAE at 30-dpi. Two-way RM ANOVAs were used with Sidak’s multiple comparisons test post-hoc. **(C)** Representative images of Western blotting for inflammasome components in BMDMs vs. C8-S cells. Cells in all groups were pre-treated with Ultrapure LPS. Groups in Lane 2 and 3 were further stimulated with nigericin and poly(dA:dT)/liposome to activate the NLRP3 and AIM2 inflammasomes, respectively. (Cells in Group1 were treated by Ultrapure LPS alone.) **(D, E)** Representative images (*D*) and quantification (*E*) of TUNEL staining of SC sections from naïve (*n*=3) and 30-dpi EAE ASC-Citrine mice (*n*=4). Two-way repeated measures (RM) ANOVAs was used. Scale bar is 75 μm. **(F, G)** Representative images *(F)* and quantification *(G)* of active caspase-3 (CC3) in SC astrocytes with and without ASC specks/strings. Evaluated from naïve (*n=7*) and at 30-dpi EAE (*n=6*) ASC-Citrine mice. Scale bar is 200 μm. Individual astrocytes were identified using the Imaris software and were quantified by CC3 puncta staining. Each datapoint represents a value from one mouse, combined from multiple experiments. Two-way RM ANOVA was used with Sidak’s multiple comparison test post-hoc. *(A, B, E, G)* \*\**p*<0.01, \*\*\**p*<0.001, \*\*\*\**p*<0.0001. Error bars denote mean ± SEM.

Next, we investigated whether astrocytes with ASC specks or strings undergo cell death during EAE. We attempted to assess general cell death by TUNEL staining. TUNEL^+^ cells were present in the periphery of the SC at 30-dpi EAE, though no TUNEL^+^ astrocytes were detected (**Fig. 6*D, E***), consistent with previous data demonstrating that astrocytes do not undergo significant cell death during EAE (28). Although normally not associated with canonical inflammasomes, we found enriched active caspase-3 (CC3) in astrocytes with active inflammasomes in both ASC-Citrine (**Fig. 6*F, G***) and *Gfap*^*Cre*^*;Asc-Citrine*^*LSL*^ (**Fig. S5*I***) mice, suggesting a possible connection between inflammasomes and caspase-3 activation in astrocytes. In summary, inflammasomes activation in astrocytes does not appear to lead to typical outcomes of inflammasome activation as seen in myeloid cells.

### AIM2 facilitates inflammasome activation in the CNS during EAE

To further assess which inflammasome is activated in astrocytes during EAE, we selected NLRP3 and AIM2 among the inflammasome sensors based on their expression by astrocytes during EAE by the Ribotag raw transcript data (**Fig. S4*A***). As *Nlrp3*^*-/-*^ mice are resistant to standard EAE, we used the Type-B EAE model to induce EAE in the *Nlrp3*^*-/-*^ background (3, 9) and confirmed that ASC-Citrine mice and *Nlrp3*^*-/-*^;ASC-Citrine developed similar disease course and severity, as expected (**Fig. S6*A***). Here, *Nlrp3*^-/-^; ASC-Citrine mice still showed comparable numbers of ASC specks with ASC-Citrine mice (**Fig. 7*A, B***), suggesting that NLRP3 is dispensable in CNS inflammasome activation during EAE. Next, we tested the AIM2 inflammasome. Congruent with recent reports (13, 29), we found AIM2 to be protective in EAE, as demonstrated by more severe disease in *Aim2*^*-/-*^ mice predominantly after disease peak, when induced with a low dose adjuvant (50 μg *Mtb* /mouse) (**Fig. 7*C, D***). The immune phenotype of *Aim2*^*-/-*^ mice in SC, iLN, and spleen at 16-dpi did not show statistically significant changes compared to WT mice, although a trend of increased T cells, microglia, and macrophages were observed, possibly reflecting the disease severity of *Aim2*^*-/-*^ mice (**Fig. S*6B-D***), together with enhanced astrogliosis (**Fig. S6E, 7E**).

**Figure 7.**
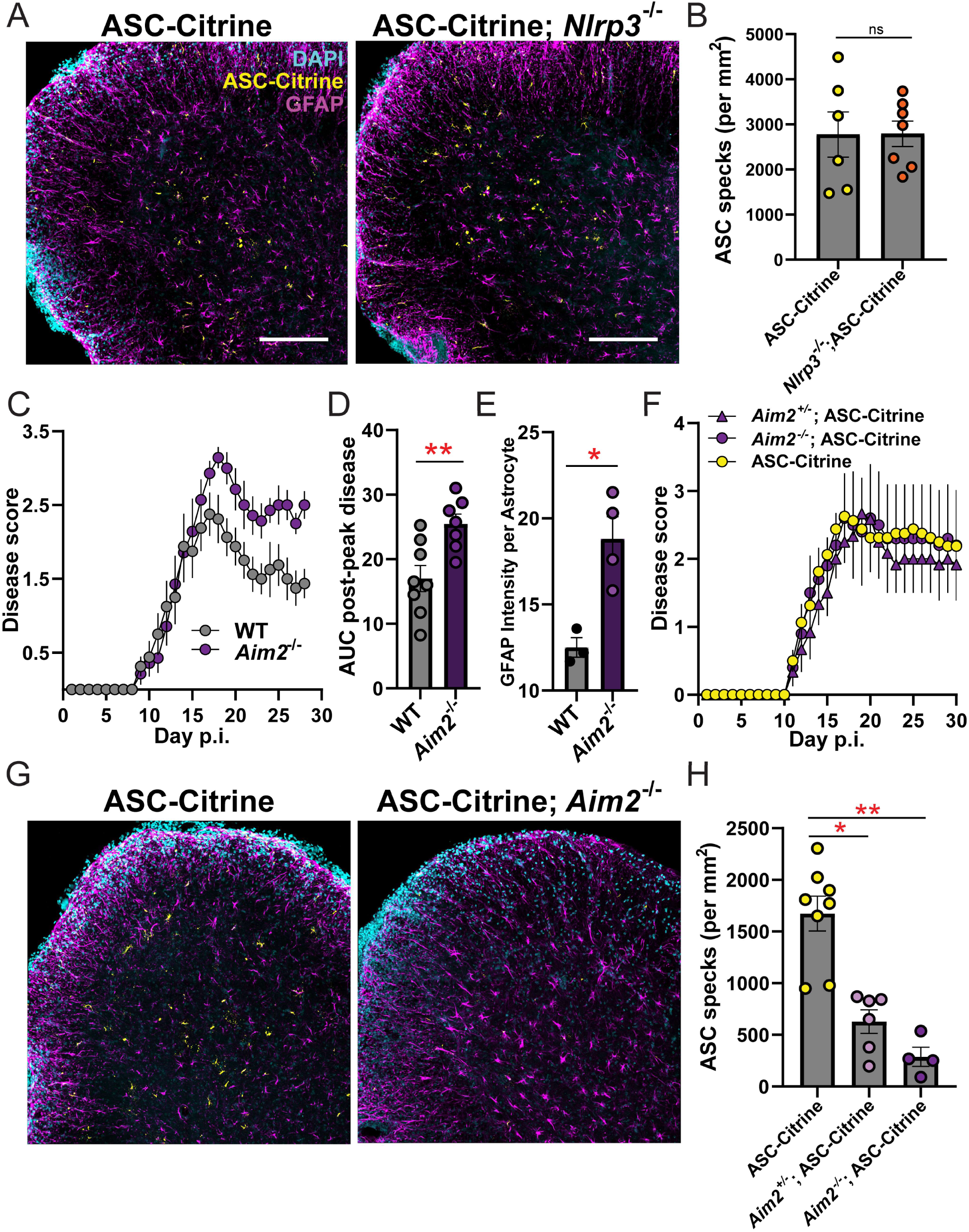
AIM2 inflammasome activation in astrocytes during EAE. **(A, B)** Comparison of ASC speck formation in indicated mouse groups. Representative images *(A)* and quantification of ASC specks *(B)* of SC sections from ASC-Citrine (*n*=6) versus *Nlrp3*^*-/-*^ ASC-Citrine (*n*=7) mice at 30-dpi Type B EAE. Mann-Whitney test was used. **(C, D)** EAE disease scores (*C*) and statistical evaluation of AUC (*D*) of WT (*n*=8) versus *Aim2*^*-/-*^ (*n*=7) mice induced with Type-A EAE with low dose *Mtb* (50 μg/mouse). Mann-Whitney test of AUC was used to analyze post-peak disease (*D*). **(E)** Quantification of GFAP intensity per cell in gray matter SC astrocytes from WT (*n*=3) versus *Aim2* ^-/-^ (n=4) mice at 30-dpi of Type-A EAE with low dose *Mtb* (50 μg/mouse). Unpaired t-test was used. **(F-H)** Type-A EAE with 200 μg *Mtb* /mouse. EAE disease score of ASC-Citrine (*n*=8), *Aim2* ^+/-^ ;ASC-Citrine (*n*=6), *Aim2* ^-/-^; ASC-Citrine (*n*=4) mice *(F)*. Representative images *(G)* and quantification of ASC specks *(H)* of SC sections at 30-dpi EAE. Scale bar is 200μm (*A, G*). One-way ANOVA was used (*p* < 0.0001) with Dunnet’s multiple comparisons test post-hoc *(H)*. Unpaired t-test was used. (*B,D,E*). Each datapoint represents a value from one mouse (*B, D, E, H*). *ns*; not significant (*p*>0.05), \**p*<0.05, \*\**p*<0.01. Error bars denote mean ± SEM.

Next, we investigated the role of the AIM2 in astrocyte inflammasome activation during EAE. To do so, we sought an EAE condition for *Aim2*^*-/-*^ mice to develop comparable EAE severity. Despite increased EAE severity in *Aim2*^*-/-*^ mice, an increased adjuvant dose (200 μg *Mtb* /mouse) elicited comparable EAE scores between WT and *Aim2*^*-/-*^ mice (**Fig. 7*F***). Here, a strong gene dosage effect of AIM2 on ASC speck formation was observed (**Fig. 7*G, H***), strongly suggesting that inflammasome activation in astrocytes *in vivo* indeed requires AIM2.

## DISCUSSION

Many studies show gene and protein expression of inflammasome components in the CNS *in vivo*, but a few have evaluated *bona fide* inflammasome activation. This study used reporter mice to detect activated inflammasomes *in situ* in a cell-type-specific manner in the CNS during EAE. Using the ASC-Citrine mice, we identified AIM2 inflammasome activation after disease peak mainly in astrocytes in the SC of EAE mice. Unexpectedly, we detected limited or no inflammasome activation in CNS-infiltrated myeloid cells or microglia.

Previous studies suggested inflammasome activation in CNS-resident cells during EAE and MS, but these studies evaluated inflammasome activation of microglia or astrocytes in tissue culture (8, 13-17). A few studies demonstrated inflammasome activation in the CNS during EAE *in situ*. One such study showed ASC specks in Iba-1^+^ cells, suggested to be microglia, in the hippocampus of EAE mice, but the investigation was not extended to astrocytes (11). Another study suggested caspase-8-mediated noncanonical NLRP3 inflammasome activation in microglia in the SC of EAE mice based on caspase-8-FLICA staining of tissue sections (30). It is possible that a small number of microglia may activate inflammasomes, although our data showed that significantly more inflammasome activation occurs at 30-dpi in astrocytes.

We and other groups have shown that the NLRP3 inflammasome is detrimental in EAE. However, recent studies (13, 29) and our results demonstrate that AIM2 can play a protective role. Chou et al. demonstrated that AIM2 suppresses EAE by promoting T_reg_ stability in an inflammasome-independent fashion (29). Similarly, the study by Ma et al. demonstrated a protective role of AIM2 through another inflammasome-independent mechanism targeting the DNA-PK-AKT3 pathway (13). Using *Aim2* ^*fl/fl*^*;Gfap*^*Cre*^ mice, Ma et al. evaluated the disease severity in Type-B EAE and found that astrocyte-specific *Aim2* depletion did not change the disease severity (13). Of note, the study did not evaluate the disease score beyond 18 dpi (around disease peak) (13). However, our data indicated numbers of ASC specks both in naïve and 16-dpi EAE mice (around disease peak) are basal, while the most significant increase was observed on 30-dpi. Additionally, Ma et al. induced Type-B EAE in *Aim2* ^*fl/fl*^*;Gfap*^*Cre*^ mice (13) but we did not. Thus, intensity of EAE induction possibly affects the involvement of AIM2 in EAE too, as we demonstrated that increased adjuvant upon induction of EAE blunts the impact of AIM2 on disease (**Fig. 7*F***). A long-term evaluation of astrocyte-specific AIM2 knockout mice and elucidating an impact of EAE induction methods will merit further understanding the protective role of the AIM2 inflammasomes in astrocytes.

Using the ASC-Citrine mouse model, two recent reports have identified ASC specks in the cerebellum during development (12) and in retinal astrocytes during ocular hypertension injury (31). Notably, one of the studies also showed the ASC specks in naïve animals (12), mirroring our finding of ASC specks in naïve SC (**Fig. *1D, F***). Further, the ASC specks in the study are AIM2-dependent, and the AIM2 inflammasome contributes to normal brain development (12). Therefore, the function of the AIM2 inflammasome in the CNS may be intrinsically different from that in peripheral myeloid cells, which are equipped to induce inflammation. For example, our study indicated that astrocytes express little *Il1b* mRNA and exhibit no marked cell death upon inflammasome activation in EAE, suggesting that inflammasome activation in astrocytes *in vivo* may have biological implications other than enhancing inflammation.

In this study, we observed ASC strings, which were unique to the CNS *in vivo*. However, some *ex vivo* studies have shown a similar ASC string-like structure. “ASC filaments” have been documented *ex vivo* with mutant ASC (32) or with ASC CARD domain blockade or deletion (33-35). An ASC isoform (ASC-c) also generates ASC filaments in human cells and appears to be expressed in mice, at least in the J774A.1 cell line (36). It is not known how astrocytes generate ASC strings, but several possibilities exist, such as potential astrocyte-specific expression of the ASC-c isoform or interaction of inflammasome components with astrocyte-specific molecules. For example, GFAP and vimentin bind together as part of the astrocyte intermediate filament network (37); and vimentin is known to interact with inflammasome components, such as caspase-1 (38). Thus, a possible physical association of GFAP to inflammasome components might explain the appearance of ASC strings in the highly ramified astrocyte.

We identified activated caspase-3 in astrocytes with ASC specks in the absence of cell death. Caspases, including caspase-3, possess critical functions outside of induction of cell death (39). Specifically, non-apoptotic caspase-3 activation is involved in the differentiation of numerous cell types, such as monocytes, neurons, and hematopoietic stem cells (39). Also, neurons, which derive from similar progenitors to astrocytes, rely on caspase-3 for dendrite and axon remodeling (40, 41) and synaptic plasticity (42). Caspase-3 activation in astrocytes is also associated with astrogliosis and not cell death (43-46). Specifically, caspase-3 activation in astrocytes is associated with cytoskeletal remodeling in a kainic-acid induced neurodegeneration model (44), reactive astrocytes following excitotoxic N-Methyl-D-aspartate (NMDA)-induced neurodegeneration (46), and GFAP cleavage in an Alzheimer’s disease model (43). Indeed, our data suggested the involvement of AIM2 inflammasome in caspase-3 activation and GFAP cleavage. The non-apoptotic activation of caspase-3 in astrocytes is inducible *in vitro*, and promotes expression of glutamate synthase and basic fibroblast growth factor-mediated by astrogliosis (45). Our study now connects inflammasome activation in astrocytes to caspase-3 activation and so warrants further study into the role of caspase-3 in astrogliosis.

The AIM2 inflammasome is activated by double-stranded DNA (dsDNA) derived not only from microbes but also from endogenous sources as a sentinel of genotoxic stress and DNA damage. The AIM2 inflammasome protects from gastrointestinal toxicity and hematopoietic failure in total-body irradiation (47), which triggers DNA double-strand breaks (47) and nuclear membrane disruption (48). Indeed, the detection of dsDNA by AIM2 is required for normal neurodevelopment during periods of proliferative stress in neurons of the CNS (12). These studies demonstrated AIM2-mediated protection from damage and even limiting inflammation, which positions the AIM2 inflammasome separately from the classical understanding of other inflammasomes, such as the NLRP3 inflammasome. Opposing outcomes of EAE severity in the absence of AIM2 versus ASC, as well as our results demonstrating a detrimental role of ASC in non-hematopoietic cells (**Fig. 2*A-C***), are intriguing; however perhaps not surprising, as ASC is the common adaptor to other inflammasomes, including the NLRP3 inflammasome, which is pathogenic during EAE. Thus, the pathogenic impact of the NLRP3 inflammasome (and perhaps the Pyrin inflammasome (49)) on EAE potentially supersedes the functions of the AIM2 inflammasome in an ASC-deficient animal.

In conclusion, our study demonstrates astrocyte AIM2 inflammasome activation without eliciting IL-1β-mediated inflammation in the late phase of EAE. This study expands our understanding of astrocytes in EAE and warrants further investigation of non-inflammatory functions of the AIM2 inflammasome in astrocytes during neuroinflammation.

## MATERIALS AND METHODS

### Animals

We used mice of the C57BL/6 genetic background of both sexes aged 8-12 weeks old, unless otherwise noted. Because we did not identify sex differences in our experiments, both male and female were equally represented in our experiments. The ASC-Citrine mice were generated and gifted by Dr. Douglas Golenbock (University of Massachusetts Medical School). The *Pycard*^-/-^ and *Nlrp3*^-/-^ mice were initially obtained from Genentech. The following mice are from The Jackson Laboratory; *Cx3cr1*^CreERT2^ (#020940) *Gfap*^Cre^ (#012886), *Asc-Citrine*^*LSL*^ (#030743), *Syn1*^Cre^ (#003966), and *Aim2* ^-/-^ (# 013144). *Gfap*^*Cre*^ and *Syn1*^Cre^ mice were used as heterozygotes. Mice for all experiments were housed in a specific pathogen-free environment. All animal experiments included in this study were approved by the Institutional Animal Care and Use Committee of Duke University.

### EAE induction and scoring

Mice were immunized with CFA/MOG emulsion in the lower back on day 0. The emulsion was prepared by mixing MOG_35-55_ peptide (United Biosystems, Cat# U104628) and complete Freund’s adjuvant (Sigma-Aldrich, Cat# F5881) with additional *Mtb* (BD Difco, Cat# 231141; 200 μg/mouse). The mice also received an intraperitoneal injection of Pertussis toxin (PTx) (200ng/mouse) (List Biological Technologies, Cat# 180) on Day 0 and 2. Unless otherwise noted, we induced EAE with this method, as “Type-A EAE” (100 μg MOG_35-55_/mouse and 200 μg *Mtb*/mouse)(9). In some experiments, “Type-B EAE” (9) was induced with CFA/MOG injection on day 0 and 7 (100 μg MOG_35-55_/mouse, 400 μg *Mtb* /mouse) and PTx (200 ng/mouse) on day 0, 2, and 7. EAE was scored as previously described (2, 3, 9).

### Preparation of frozen tissue sections and staining with antibodies

Animals were lethally anesthetized with 100 mg/kg of Nembutal administered intraperitoneally and transcardially perfused with PBS and subsequently 4% paraformaldehyde (PFA)(Sigma-Aldrich, Cat# 158127). SC and iLNs were harvested and fixed for 24 hours at 4°C in 4% PFA. All tissues were cryoprotected in 30% sucrose in water for an additional 24 hours before embedding and freezing in Tissue-Tek O.C.T. compound (Sakura, Cat# 4583) on dry ice. Tissues were sectioned using a Cryostar NX50 (Thermo Fisher Scientific) at a thickness of 25 µm and rendered as floating sections. Sections were permeabilized with 0.25 % Triton X100 (Amresco, Cat# 0694-1L) and blocked using 2% bovine serum albumin (GeminiBio / Cat# 700-101P). After antibody staining, sections were mounted onto slides with ProLong™ Gold Antifade Mountant (Invitrogen, Cat# P36931) or ProLong™ Gold Antifade Mountant with DAPI (Invitrogen, Cat# P36930). TUNEL staining was also performed with 25 μm thick floating tissue sections with the CF 640R TUNEL Assay Apoptosis Detection Kit (Biotium, Cat# 30074). Antibodies used for staining are indicated in **Supplementary Table 1**.

### Preparation of tissue slice culture and anti-ASC immunostaining

Spleens were dissected out of mice, and live spleen slices were prepared with a vibratome (Precisionary Compresstome) using 4% low melt agarose as described by manufacturer’s protocol. Slices in complete RPMI were kept as floating sections and treated with 100 ng/mL Ultrapure LPS (Invivogen, Cat# tlrl-3pelps) for 2 hours followed by 5 μM nigericin (Sigma-Aldrich, Cat# N7143) for additional 1 hour. After fixation and permeabilization with ice cold methanol for 15 minutes, slices were blocked with 2% BSA in PBS for 1 hour at RT. Immunofluorescence staining was performed with antibodies listed in **Table S1**.

### Immunofluorescence microscopy and image analyses

All slides were imaged on the Zeiss 710 Inverted Laser Scanning Confocal Microscope (Duke University Light Microscopy Core Facility) at full 25 µm depth as z-stacks. For quantifications, 2 sections per animal were imaged, and a 2×2 grid tile scan was performed using the 20x objective centered on the ventral horn of the SC, totaling 8 fields per mouse. Following quantification, these replicates were averaged to generate a single *n* for statistical analyses. Semi-automated quantification was conducted using the Imaris for Neuroscientists Cell Imaging Software ver. 9.3.0. (Bitplane) unless otherwise indicated. Briefly, the Surfaces tool was used to identify either ASC specks/strings or cells, and intensity thresholds of counterstain signals within the surfaces were used to quantify the desired characteristics. ASC specks and ASC strings counts in **Fig. 1** were manually enumerated using the LSM Browser software (Zeiss).

### Bone Marrow Chimeras

Recipient mice (CD45.2; 6-8 weeks old) were lethally irradiated with 900 rad (XRAD 320 X-Ray irradiator) and adoptively transferred with 10^6^ CD45.1 donor bone marrow (BM) cells. Mice were supported with water containing sulfamethoxazole and trimethoprim for 1 week following irradiation. At 6 weeks post-adoptive transfer, the donor cell reconstitution was confirmed by differential expression of congenic markers in peripheral blood using the BD FACSCanto II (BD Biosciences).

### Tamoxifen pulse for selective microglial labelling

This procedure was adapted from previous work (50). Briefly, mice with *Cx3cr1*^*CreERT2*^ (6-8 weeks old) received an intraperitoneal injection of 75 mg/kg tamoxifen dissolved in corn oil. The second injection with the same formula was administered two days later. Mice were kept for six weeks after the last tamoxifen administration for six weeks before EAE induction.

### Astrocyte isolation and RT-qPCR

Astrocytes were isolated as previously described (51) with a few modifications. Briefly, mouse SC were minced into ∼1mm^2^ pieces and digested using the Papain Dissociation System (Worthington Biochemical Corporation, Cat# LK003153). Isolated cells were negatively selected tandemly first with Myelin Removal Beads II (Miltenyi Biotec, Cat# 130-096-731) and second with anti-CD11b MicroBeads (Miltenyi Biotec, Cat# 130-049-601) to collect the flowthrough fraction. To enrich astrocytes, the flowthrough cells were first treated with FcBlock (Miltenyi Biotec, Cat# 130-092-575), then positively selected using anti-ACSA-2 MicroBeads (Miltenyi Biotec, Cat# 130-097-678). Total RNA was prepared using TRI Reagent (Millipore-Sigma, Cat# 93289) and reverse-transcribed using qScript cDNA Mix (Quantabio, Cat# 950048) to obtain cDNA. RT-qPCR assays were performed with SYBR FAST qPCR Master Mix (Kapa Biosystems, Cat# KK4602), using primers indicated in **Supplementary Table 2**. Expression levels of target genes relative to an internal control, *Actb*, were calculated using the *– ΔΔCt* method (52).

### Cell culture and Western blotting analysis

BMDMs were generated by culturing total BM cells for 7 days in complete RPMI supplemented with 10 ng/mL recombinant mouse M-CSF (BioLegend, Cat# 576406M-CSF). C8-S cells (ATCC, Cat# CRL-2535) were cultured in complete DMEM. One day before stimulation, BMDMs and C8-S cells were seeded into poly-L-lysine-coated 12-well plates (1.5 × 10^6^ cells/well). Then, cells were pre-treated with 100 ng/mL Ultrapure LPS (Invivogen, Cat# tlrl-3pelps) in serum-free Opti-MEM medium (Thermo Fisher Scientific, Cat# 11058021) for 2 hours and stimulated with 5 μM nigericin (Sigma-Aldrich, Cat# N7143) or poly(dA:dT) (InvivoGen, Cat# tlrl-patn) for 4 hours to activate the NLRP3 and AIM2 inflammasomes, respectively. Poly(dA:dT) was used as a complex with Lipofectamine 2000 (InvitroGen, Cat# 11668019) at a concentration of 0.5 μg/mL. Culture supernatants and cell lysates (in RIPA buffer) were harvested and, total protein concentrations were quantified using the BCA Protein Assay Kit (ThermoFisher Scientific, Cat# 23227). The same amount of total protein was used across all samples for SDS-PAGE gel separation. Western blotting (WB) was performed with indicated antibodies (**Supplementary Table 1**), and the result was imaged using the GeneGnome Chemiluminescence System (Syngene) and the Genesys (Syngene) software. Band intensity was quantified using the Genetools (Syngene) Software. Lysate β-actin band intensity was used to normalize both the lysate and supernatant band intensity.

### Statistical Analyses

Area under the curve (AUC) values were used to statistically analyze EAE scoring data. The Mann-Whitney U-test was used to compare between two groups, unless otherwise indicated. Pre-peak and post-peak AUC were defined by identifying the dpi at which disease score ceased to increase or began to decrease for both groups in a single experiment. All other analyses, where indicated in the figure legend, were performed using either the Mann-Whitney U-test, a One-Factor ANOVA, or a Two-Factor ANOVA. Post-hoc testing was performed only if ANOVA reached significance on interaction term (*p < 0*.*1*). The Dunnett’s Multiple Comparisons and the Sidak’s multiple comparisons were conducted post-hoc where indicated in the figure legend. All statistical analyses were performed using the Graphpad Prism 8 software.

## Author Contributions

W.E.B., M.E.D., and M.L.S. designed research; W.E.B., M.E.D., M.I., T.N., K.N., N.A.L., and N.A. performed experiments, W.E.B. M.E.D., and M.L.S. analyzed data; W.E.B. and M.L.S wrote the manuscript; and M.E.D., M.I., K.N., N.A.L., N.A. and E.A.M. edited the manuscript.

## Data Availability

All study data are included in the article and/or *SI Appendix*.

## Acknowledgements

We appreciate Dr. Golenbock for his generous gift of a mutant mouse strain and for sharing unpublished data. We also appreciate Dr. Cagla Eroglu and Maria Pia Rodriguez Salazar for their advice on handling astrocytes. We appreciate Dr. Ryan Finethy for his advice on inflammasome immunostaining protocols. We also appreciate Tomoko Kadota and Amesha Crudup for their help in mouse maintenance. This study was funded to M.L.S. by National Multiple Sclerosis Society (NMSS) Research Grant (RG 4536B2/1), NIH (R01-NS120417, R01-AI088100), and the Chancellor’s Discovery Program Research Fund at Duke University School of Medicine.

## Footnotes

<u>↵</u>^1^To whom correspondence may be addressed.

**AIM2**

## Figure Legends

**Figure S1.**
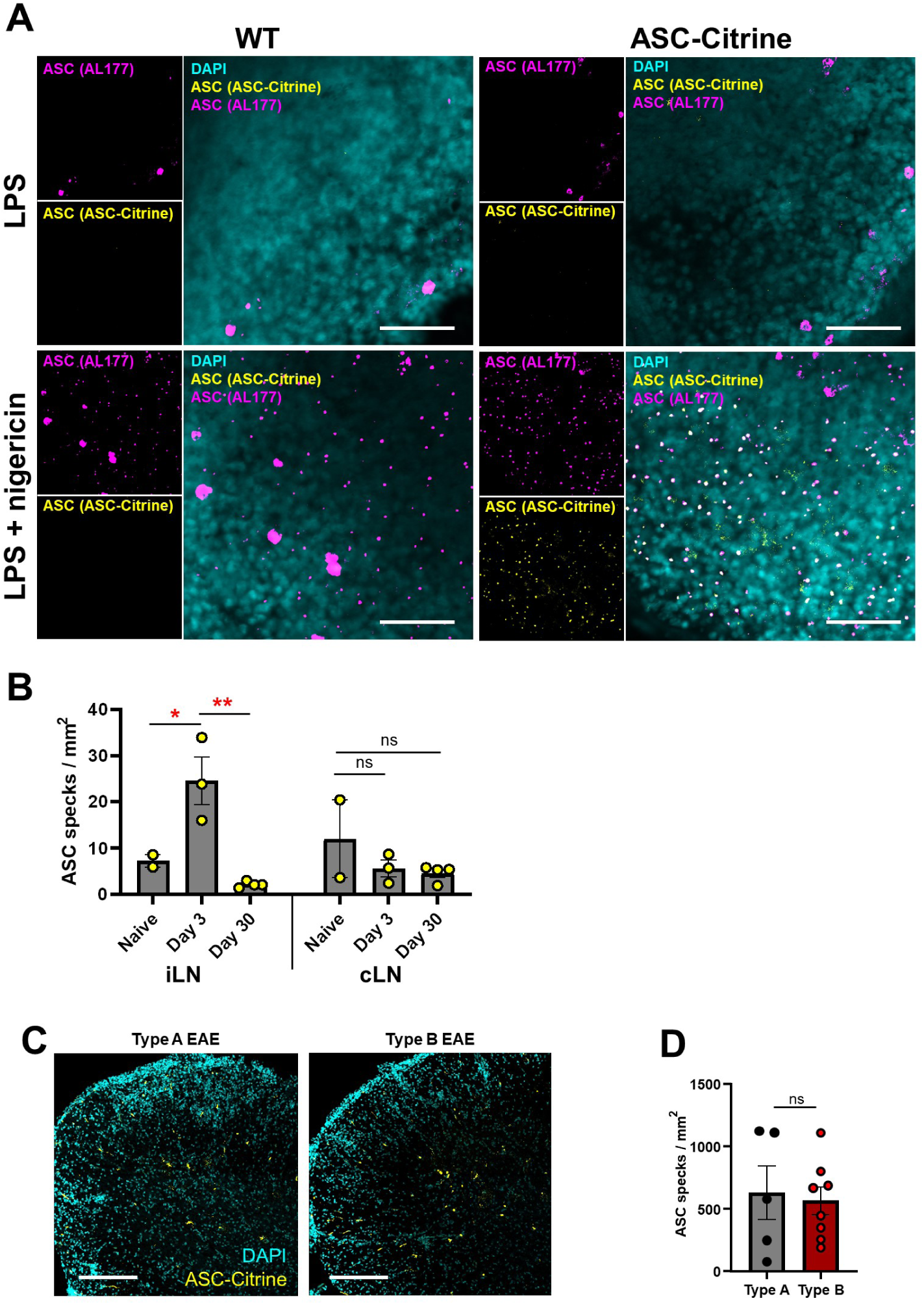
Validation of ASC-Citrine system in tissue imaging. **(A)** Representative images of ASC speck formation detected with ASC-Citrine and ASC antibody signals. Live spleen tissue culture slices from naïve WT and ASC-Citrine mice were used with NLRP3 inflammasome stimulation. Scale bar is 50 µm. **(B)** Quantification of ASC specks in the iLNs and cLNs of ASC-Citrine mice during EAE. Each datapoint represents a value of an average value from two cross-sections of LNs (25 μm thickness) from one mouse. One-way ANOVA, *p*=0.0021 (iLN), *p*=0.3235 (cLN), with Dunnett’s multiple comparisons test. **(C, D)** Comparison of ASC speck images and numbers in SC between Type-A and Type-B EAE. Representative images (C) and quantification (D) of ASC specks in the SC of ASC-Citrine mice at 30-dpi for Type A (*n=5*) and Type B (*n=8*) EAE. Each datapoint represents a value from one mouse. Mann-Whitney test was used. Scale bar is 200 μm. (*B,D*) *ns*; not significant (*p*>0.05), \**p*<0.05, \*\**p*<0.01. Error bars denote mean ± SEM.

**Figure S2.**
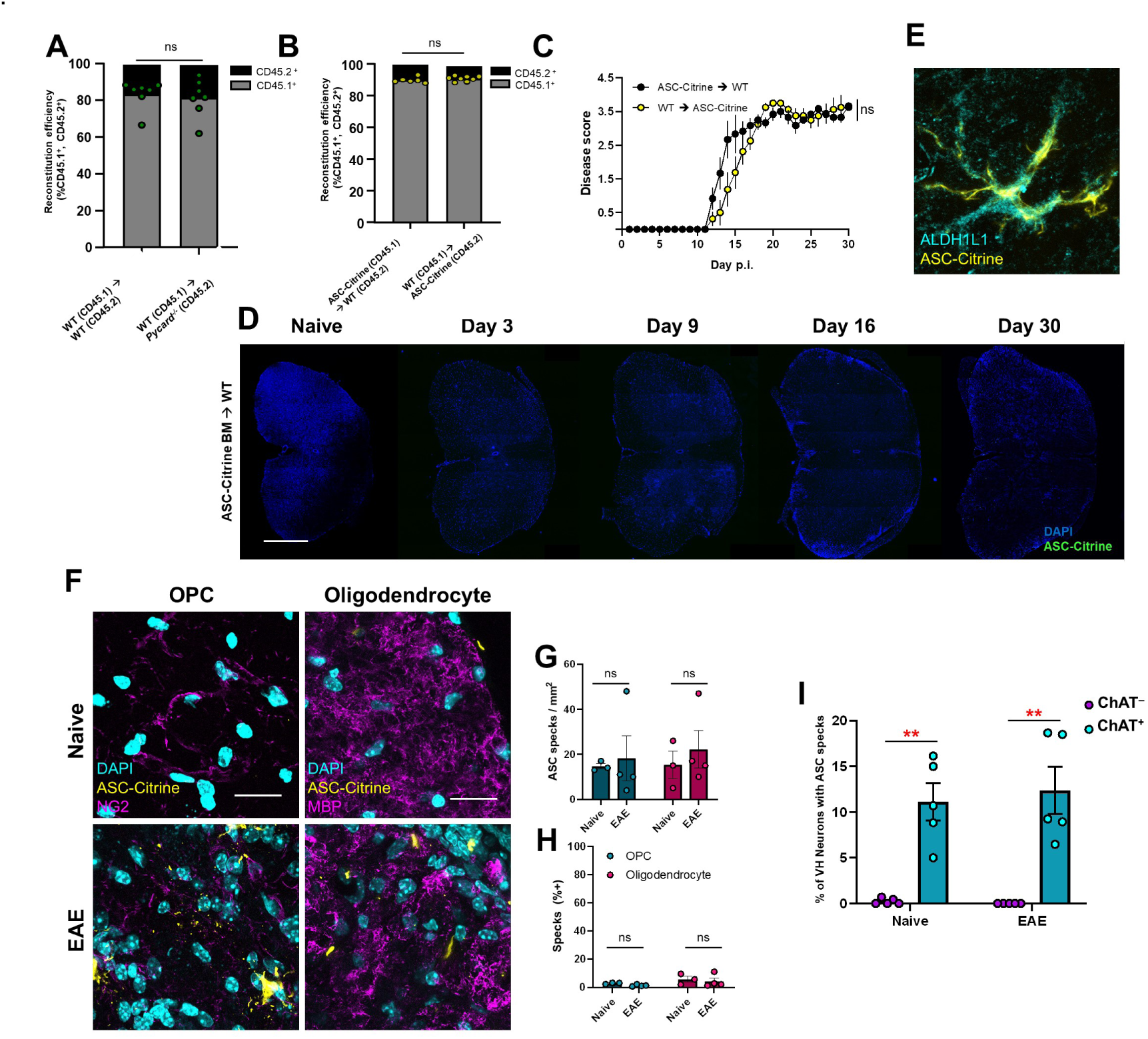
Validation of Bone Marrow Chimeras. BM chimera were created by transferring WT BM cells to irradiated WT or *Pycard*^*-/-*^ receipients (n=*7* for each group). Reconstitution efficiency of BM chimeras determined by flow cytometry, quantified as % of total CD45^+^ cells in peripheral blood for congenic markers of CD45.1 (donor) or CD45.2 (recipient). Each datapoint represents a value from one mouse. Mann-Whitney test used. **(B, C)** BM chimera were created by transferring ASC-Citrine BM cells irradiated WT recipients (ASC-Citrine → WT, *n*=6) and vice versa (WT → ASC-Citrine mice, *n*=8). Reconstitution efficiency (*B*) and EAE disease score (*C*) of indicated BM chimera Mann-Whitney test of total AUC for disease. **(D)** Representative images of SC from ASC-Citrine → WT mice at indicated time points during EAE. No apparent ASC specks were observed. Scale bar is 500 μm. **(E)** Representative image of ALDH1L1 counterstaining of astrocytes in ASC-Citrine mice at 30-dpi EAE. Scale bar is 10 μm. **(F)** Representative images of ASC specks and strings counter-stained with antibodies against NG2 (for OPCs) and MBP (for mature oligodendrocytes) in SC from naïve versus 30-dpi EAE ASC-Citrine mice. Scale bar is 20 μm. **(G)** Quantification of ASC specks in OPCs and mature oligodendrocytes of SC from naïve versus 30-dpi EAE ASC-Citrine mice. Each datapoint represents a value from one mouse. Two-way repeated measures (RM) ANOVA was used (main effect of cell type: ^ns^*p*<0.7807). **(H)** Percentages of ASC specks detected in OPC or mature oligodendrocytes out of total ASC specks per section, indicating relative contribution of OPCs and oligodendrocyte ASC specks to total number of ASC specks in L5 spinal cord at 30-dpi EAE. (*A,B,C,G,H*) *ns*; not significant (*p*>0.05). Error bars denote mean ± SEM. **(I)** Percentage of ChAT^+^ and ChAT^−^ VH neurons containing ASC specks in SC from naïve vs. 30-dpi EAE ASC-Citrine mice. Each datapoint represents a value from one mouse (n=5). Two-way RM ANOVA was used (main effect of cell type: p<0.001) with Sidak’s multiple comparisons test post hoc. **p<0.01.

**Figure S3.**
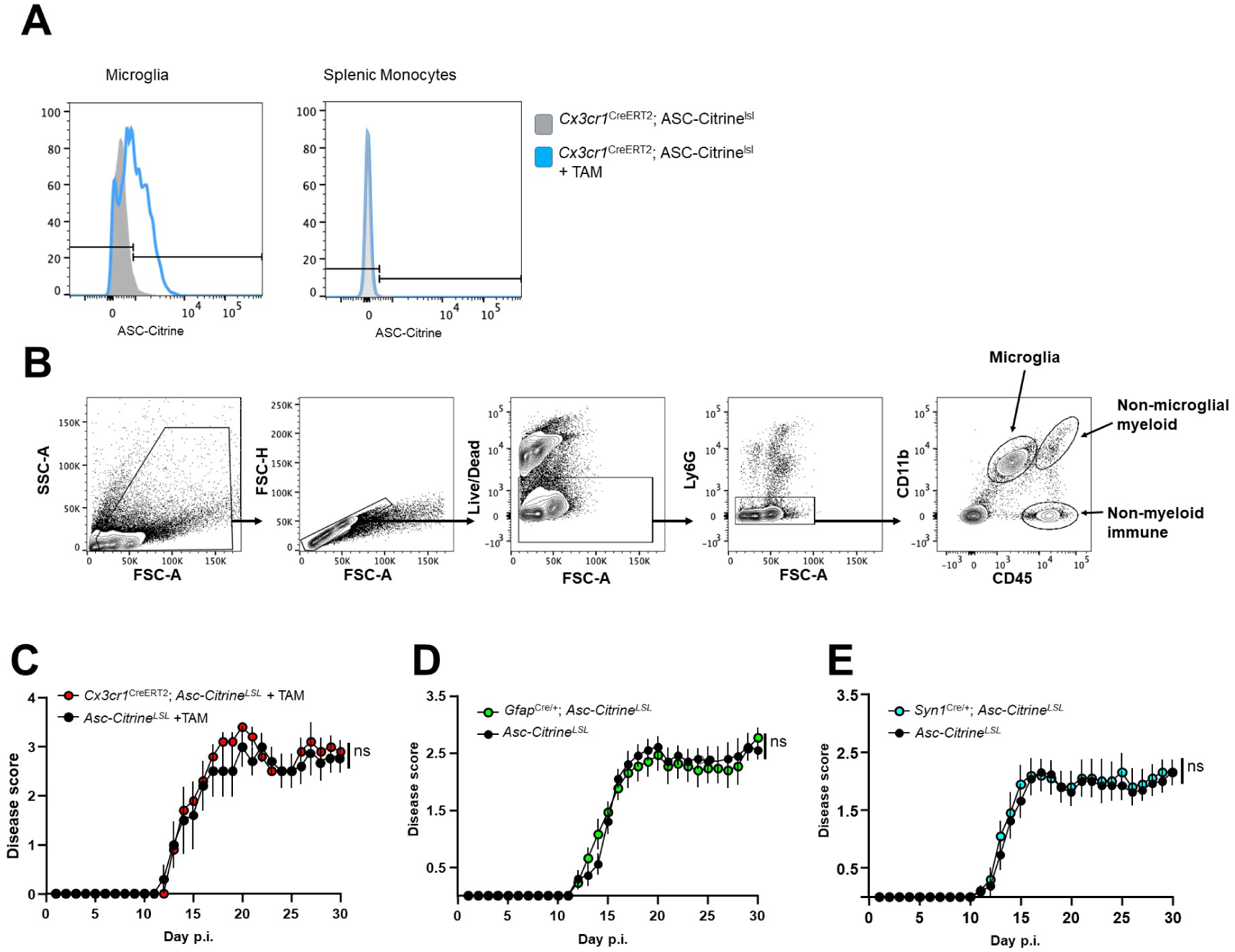
Validation of EAE mice with cell type-specific ASC-Citrine expression. **(A)** Tamoxifen mediated expression of ASC-citrine reporter expression in microglia and splenic monocytes in *Cx3cr1*^*CreERT2*^*;Asc-Citrine*^*LSL*^ with or without tamoxifen (TAM) treatment. **(B)** Flow cytometry gating strategy for identifying microglia. **(C)** EAE disease score of *Cx3cr1*^CreERT2^*;Asc-Citrine*^*LSL*^ (*n=5)* vs. *Cx3cr1*^CreERT2^*;Asc-Citrine*^*LSL*^ (*n=5)*. Both groups were treated with TAM. **(D, E)** EAE disease score of *Asc-Citrine*^*LSL*^ (*n=10*) vs. *Gfap*^Cre^*;Asc-Citrine*^*LSL*^ (*n=13*) (*D*) and *Asc-Citrine*^*LSL*^ (*n=13*) vs. *Syn1*^Cre^*;Asc-Citrine*^*LSL*^ (*n=10*) (E). Mann-Whitney test of total AUC of disease score was used (*C,D,E*). *ns*; not significant (*p*>0.05). Error bars denote mean ± SEM.

**Figure S4.**
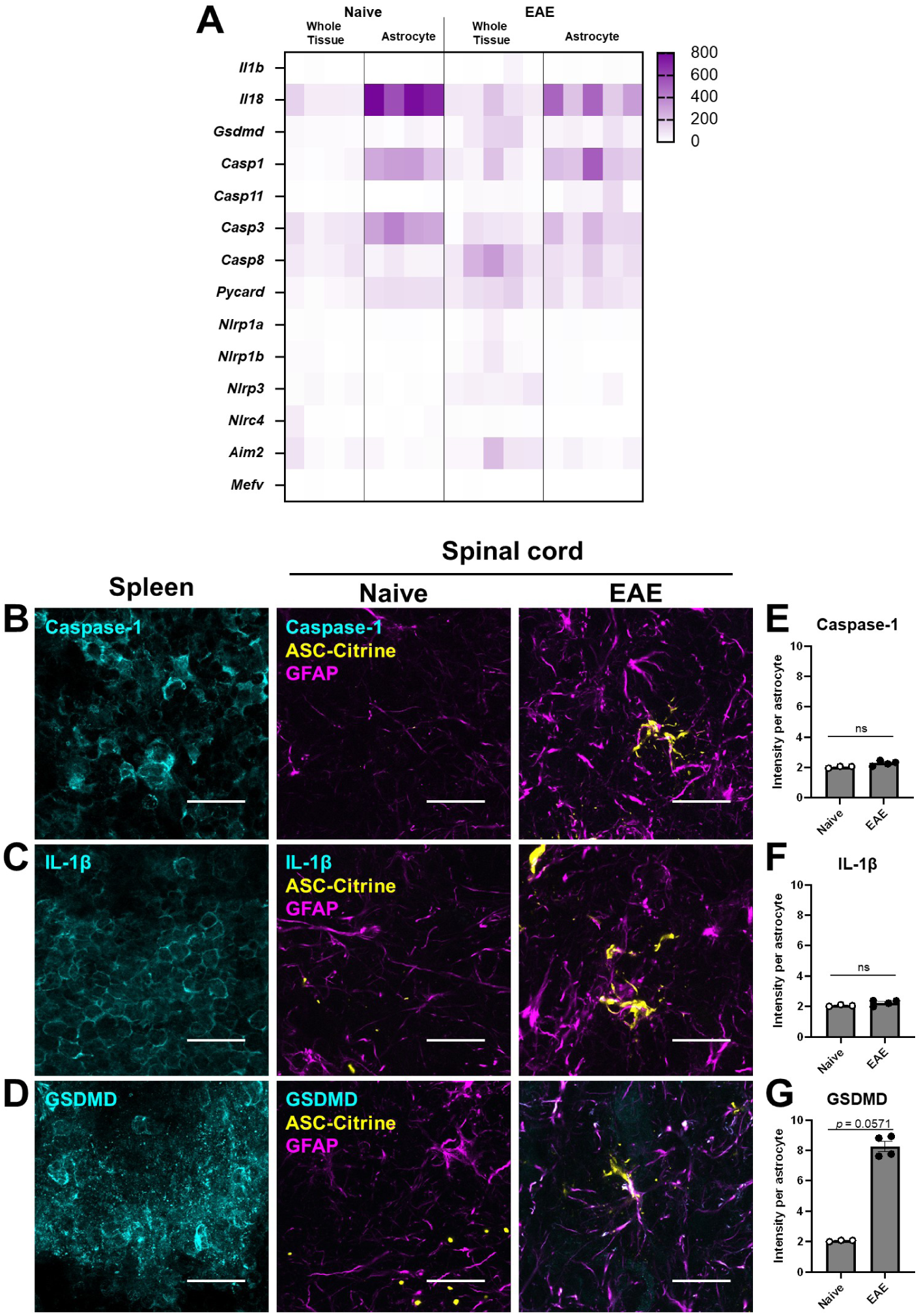
Expression of inflammasome components and cell death markers in astrocytes during EAE. **(A)** Gene-set enrichment analysis of inflammasome-associated genes in bulk SC lysates and astrocytes (with astrocyte-specific Ribotag-HA enriched RNA) in naïve and 30-dpi EAE mice. Data represented as raw transcript counts derived from publicly available data (GEO Accession #: GSE100329). **(B-G)** Representative images (*B-D*) and quantification (*E-F*) of caspase-1 (*B, E*), IL-1β (*C, F*), and GSDMD (*D, G*) expression in spleen and SC astrocytes from naïve versus 30-dpi EAE ASC-Citrine mice. Scale bar is 20 μm. Each datapoint represents a value from one mouse. Individual astrocytes were identified using the Imaris software and the mean intensity per cell was quantified for caspase-1 (*E*), IL-1β (*F*) and GSDMD (*G*). Mann-Whitney test was used. (E*-G*) *ns*; not significant (*p*>0.05). Error bars denote mean ± SEM.

**Figure S5.**
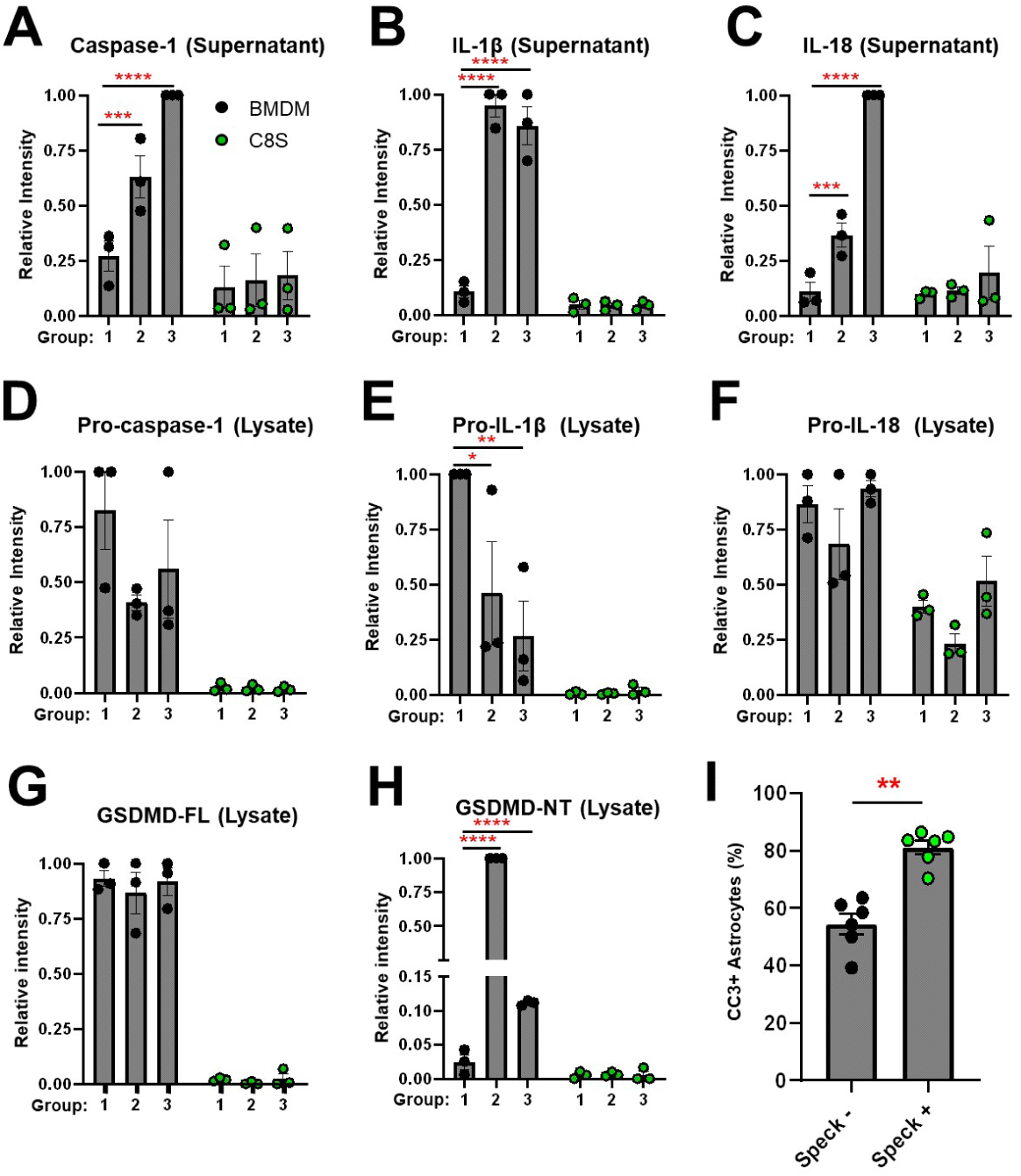
Expression of inflammasome components in C8-S astrocyte cell line. **(A-C)** WB quantitative evaluation of culture supernatant samples of mature caspase-1 (*A*), IL-1β (*B*), and IL-18 (*C*). **(D-H)** WB quantitative evaluation of cell lysate samples of pro-caspase-1 (*D*), pro-IL-1β (*E*), pro-IL-18 (*F*), GSDMD-FL (*G*), and GSDMD-NT (*H*). In (*A-H*), all groups were stimulated with Ultrapure LPS, and Groups 2 and 3 were further stimulated with nigericin and poly(dA:dT)/liposome to activate the NLRP3 and AIM2 inflammasomes, respectively. (Thus, Group 1 has Ultrapure LPS treatment alone.) Each datapoint is obtained from one independent experiment. Two-way RM ANOVA was used with Sidak’s multiple comparison test post-hoc. **(I)** Quantification of active caspase-3 (CC3) in spinal cord astrocytes with and without ASC specks/strings in *Gfap*^Cre^*;Asc-Citrine*^*LSL*^ (*n*=6) mice at 30-dpi EAE. Each datapoint represents a value from one mouse. Individual astrocytes were identified using the Imaris software and were quantified by CC3 puncta staining. Mann-Whitney test was used. (*A-I*) \*\**p*<0.01, \*\**p*<0.005, \*\*\**p*<0.001, \*\*\*\**p*<0.0001. Error bars denote mean ± SEM.

**Figure S6.**
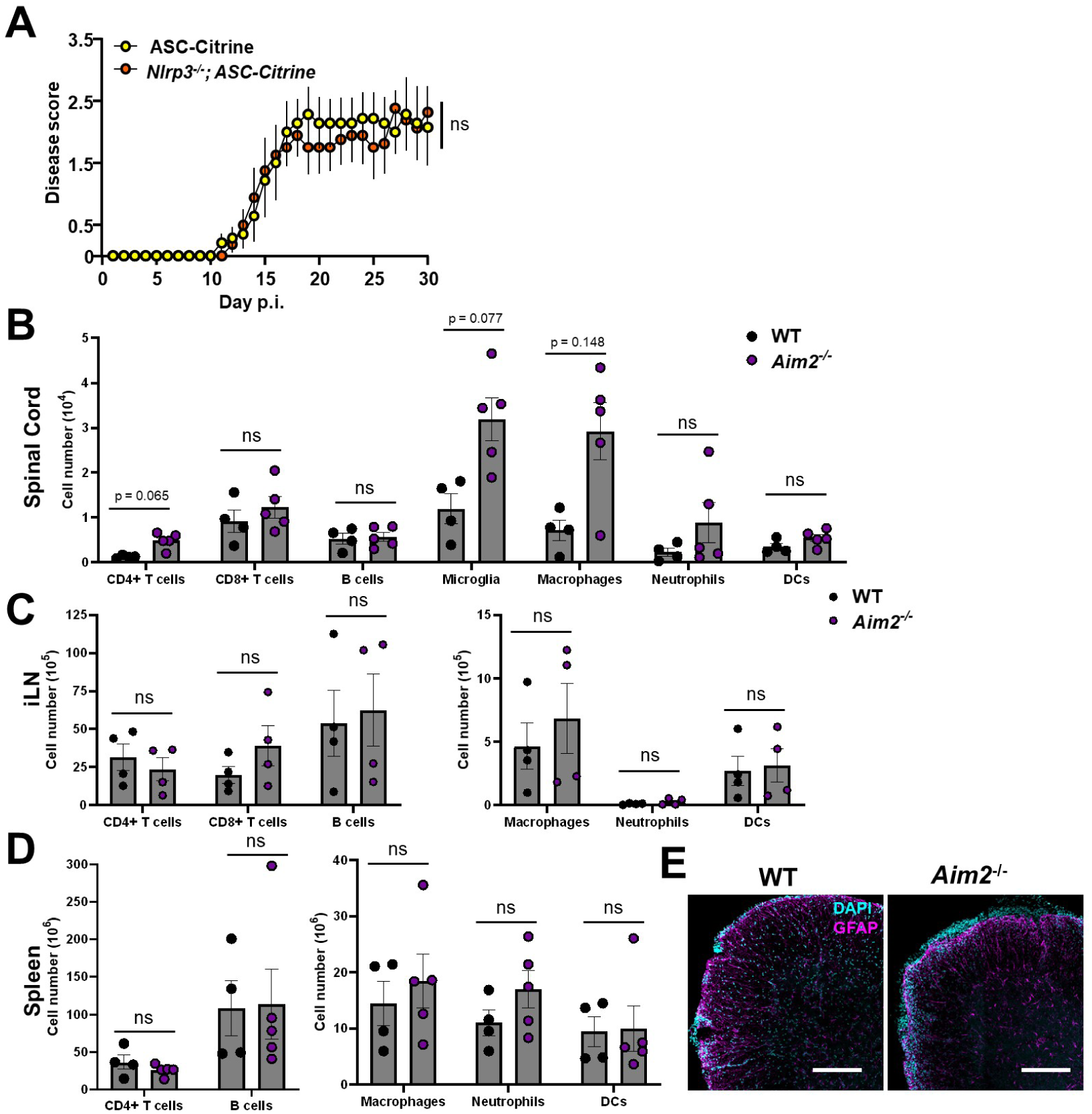
Validation of EAE phenotype of *Nlrp3*^-/-^;ASC-Citrine mice and immune phenotype of *Aim2*^*-/-*^ mice with EAE. **(A)** EAE disease score of ASC-Citrine (*n=7*) vs. *Nlrp3*^-/-^;ASC-Citrine (*n=8*) mice with Type B-EAE. Mann-Whitney test of total AUC for disease score was used. **(B–D)** Leukocyte counts in SC (*B*), iLN (*C*) and spleen (D) at 16-dpi EAE in WT vs. *Aim2*^-/-^ mice induced with Type-A EAE. One datapoint represents a value from one mouse. Two-way RM ANOVA was used with Sidak’s multiple comparisons test post hoc. **(E)** Representative image of GFAP staining in SC from WT versus *Aim2* ^-/-^ mice at 30-dpi of EAE. Scale bar is 200 μm. (*A-D*) *ns*; not significant (*p*>0.05). Error bars denote mean ± SEM.

**Table S1.**
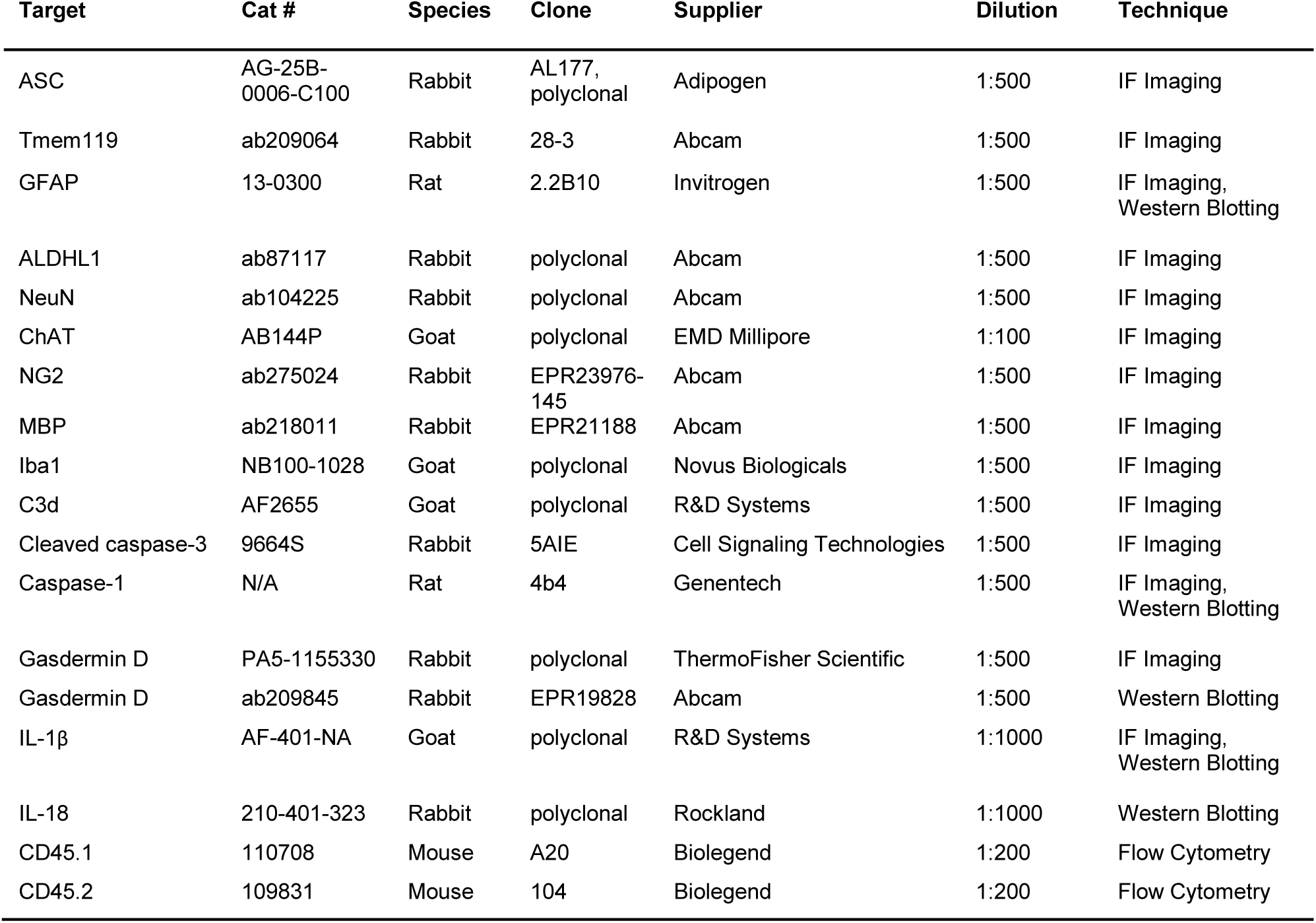
Reagent information for antibody-based techniques.

**Table S2.**
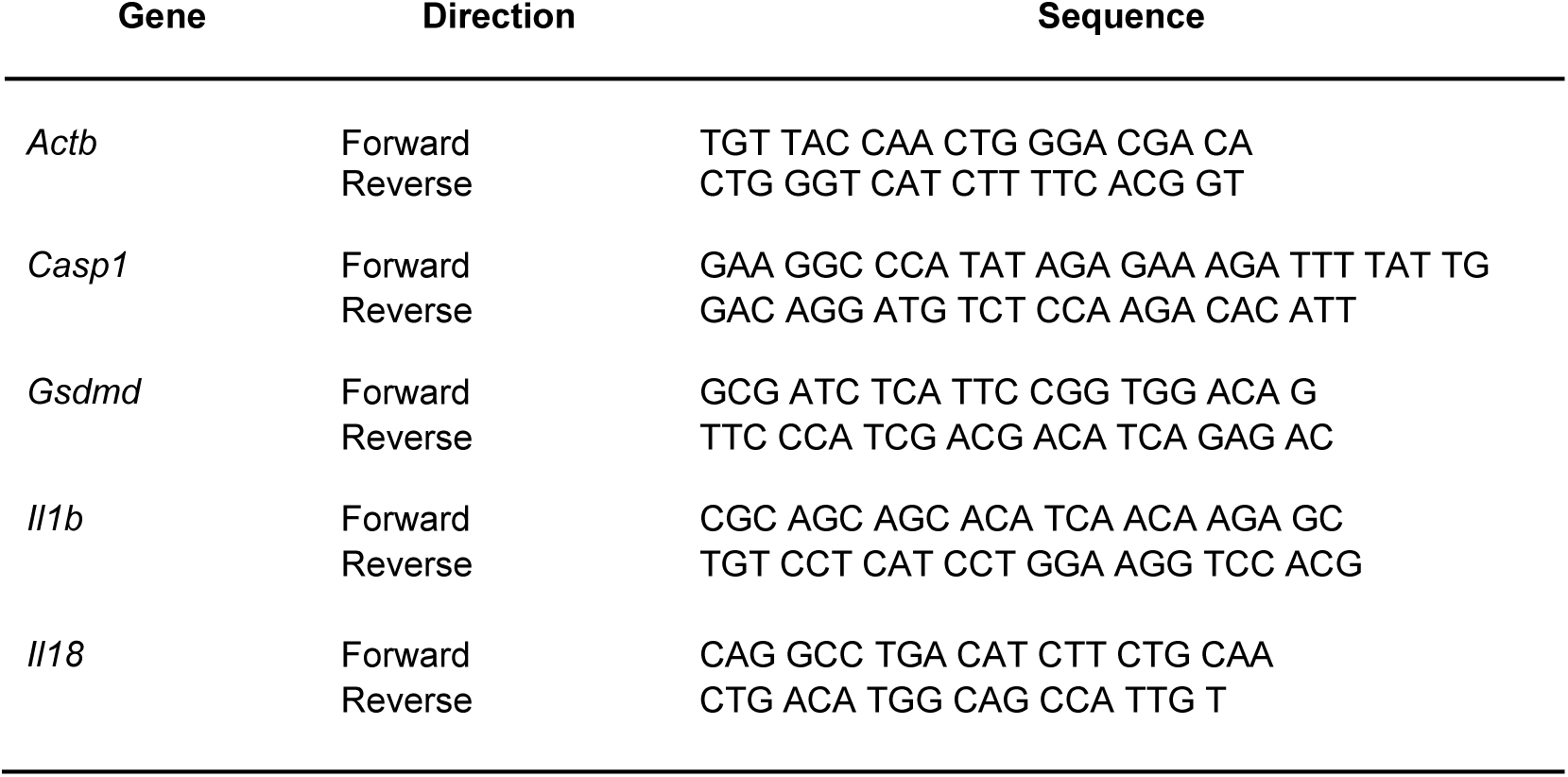
Sequences of primers used for RT-qPCR assays.

## REFERENCES

1. P. Broz, V. M. Dixit, Inflammasomes: mechanism of assembly, regulation and signalling. Nat Rev Imunol 16, 407–420 (2016).

2. M. Inoue, K. L. Williams, M. D. Gunn, M. L. Shinohara, NLRP3 inflammasome induces chemotactic immune cell migration to the CNS in experimental autoimmune encephalomyelitis. Proc Natl Acad Sci U S A 109, 10480–10485 (2012).

3. M. Inoue et al., Interferon-beta therapy against EAE is effective only when development of the disease depends on the NLRP3 inflammasome. Sci Signal 5, ra38 (2012).

4. D. Gris et al., NLRP3 plays a critical role in the development of experimental autoimmune encephalomyelitis by mediating Th1 and Th17 responses. J Imunol 185, 974–981 (2010).

5. W. Barclay, M. L. Shinohara, Inflammasome activation in multiple sclerosis and experimental autoimmune encephalomyelitis (EAE). Brain Pathol 27, 213–219 (2017).

6. L. Vidmar et al., Multiple Sclerosis patients carry an increased burden of exceedingly rare genetic variants in the inflammasome regulatory genes. Sci Rep 9, 9171 (2019).

7. S. Malhotra et al., NLRP3 polymorphisms and response to interferon-beta in multiple sclerosis patients. Mult Scler 24, 1507–1510 (2018).

8. S. Malhotra et al., NLRP3 inflammasome as prognostic factor and therapeutic target in primary progressive multiple sclerosis patients. Brain 10.1093/brain/awaa084 (2020).

9. M. Inoue et al., An interferon-beta-resistant and NLRP3 inflammasome-independent subtype of EAE with neuronal damage. Nat Neurosci 19, 1599–1609 (2016).

10. M. T. Heneka et al., NLRP3 is activated in Alzheimer’s disease and contributes to pathology in APP/PS1 mice. Nature 493, 674–678 (2013).

11. B. Hou et al., Inhibition of the NLRP3-inflammasome prevents cognitive deficits in experimental autoimmune encephalomyelitis mice via the alteration of astrocyte phenotype. Cell Death Dis 11, 377 (2020).

12. C. R. Lammert et al., AIM2 inflammasome surveillance of DNA damage shapes neurodevelopment. Nature 580, 647–652 (2020).

13. C. Ma et al., AIM2 controls microglial inflammation to prevent experimental autoimmune encephalomyelitis. J Exp Med 218 (2021).

14. B. A. McKenzie et al., Activation of the executioner caspases-3 and -7 promotes microglial pyroptosis in models of multiple sclerosis. J Neuroinflamation 17, 253 (2020).

15. B. A. McKenzie et al., Caspase-1 inhibition prevents glial inflammasome activation and pyroptosis in models of multiple sclerosis. Proc Natl Acad Sci U S A 115, E6065–E6074 (2018).

16. T. Zeis et al., Metabolic gene expression changes in astrocytes in Multiple Sclerosis cerebral cortex are indicative of immune-mediated signaling. Brain Behav Imun 48, 313–325 (2015).

17. L. Freeman et al., NLR members NLRC4 and NLRP3 mediate sterile inflammasome activation in microglia and astrocytes. J Exp Med 214, 1351–1370 (2017).

18. T. C. Tzeng et al., A Fluorescent Reporter Mouse for Inflammasome Assembly Demonstrates an Important Role for Cell-Bound and Free ASC Specks during In Vivo Infection. Cell Rep 16, 571–582 (2016).

19. G. C. Furtado et al., Swift entry of myelin-specific T lymphocytes into the central nervous system in spontaneous autoimmune encephalomyelitis. J Imunol 181, 4648–4655 (2008).

20. M. van Zwam et al., Surgical excision of CNS-draining lymph nodes reduces relapse severity in chronic-relapsing experimental autoimmune encephalomyelitis. J Pathol 217, 543–551 (2009).

21. M. J. C. Jordao et al., Single-cell profiling identifies myeloid cell subsets with distinct fates during neuroinflammation. Science 363 (2019).

22. R. Brambilla, The contribution of astrocytes to the neuroinflammatory response in multiple sclerosis and experimental autoimmune encephalomyelitis. Acta Neuropathol 137, 757–783 (2019).

23. G. Ponath, C. Park, D. Pitt, The Role of Astrocytes in Multiple Sclerosis. Front Imunol 9, 217 (2018).

24. M. Pekny, M. Pekna, Astrocyte reactivity and reactive astrogliosis: costs and benefits. Physiol Rev 94, 1077–1098 (2014).

25. M. V. Sofroniew, H. V. Vinters, Astrocytes: biology and pathology. Acta Neuropathol 119, 7–35 (2010).

26. S. A. Liddelow et al., Neurotoxic reactive astrocytes are induced by activated microglia. Nature 541, 481–487 (2017).

27. N. Itoh et al., Cell-specific and region-specific transcriptomics in the multiple sclerosis model: Focus on astrocytes. Proc Natl Acad Sci U S A 115, E302–E309 (2018).

28. F. Haroon et al., Gp130-dependent astrocytic survival is critical for the control of autoimmune central nervous system inflammation. J Imunol 186, 6521–6531 (2011).

29. W. C. Chou et al., AIM2 in regulatory T cells restrains autoimmune diseases. Nature 10.1038/s41586-021-03231-w (2021).

30. C. J. Zhang et al., TLR-stimulated IRAKM activates caspase-8 inflammasome in microglia and promotes neuroinflammation. J Clin Invest 128, 5399–5412 (2018).

31. A. Pronin et al., Inflammasome Activation Induces Pyroptosis in the Retina Exposed to Ocular Hypertension Injury. Front Mol Neurosci 12, 36 (2019).

32. A. C. Sahillioglu, F. Sumbul, N. Ozoren, T. Haliloglu, Structural and dynamics aspects of ASC speck assembly. Structure 22, 1722–1734 (2014).

33. M. Moriya et al., Role of charged and hydrophobic residues in the oligomerization of the PYRIN domain of ASC. Biochemistry 44, 575–583 (2005).

34. A. Lu et al., Unified polymerization mechanism for the assembly of ASC-dependent inflammasomes. Cell 156, 1193–1206 (2014).

35. F. I. Schmidt et al., A single domain antibody fragment that recognizes the adaptor ASC defines the role of ASC domains in inflammasome assembly. J Exp Med 213, 771–790 (2016).

36. N. B. Bryan et al., Differential splicing of the apoptosis-associated speck like protein containing a caspase recruitment domain (ASC) regulates inflammasomes. Journal of inflamation (London, England) 7, 23 (2010).

37. A. Viedma-Poyatos, Y. de Pablo, M. Pekny, D. Perez-Sala, The cysteine residue of glial fibrillary acidic protein is a critical target for lipoxidation and required for efficient network organization. Free Radic Biol Med 120, 380–394 (2018).

38. G. dos Santos et al., Vimentin regulates activation of the NLRP3 inflammasome. Nat Comun 6, 6574 (2015).

39. S. Shalini, L. Dorstyn, S. Dawar, S. Kumar, Old, new and emerging functions of caspases. Cell Death Differ 22, 526–539 (2015).

40. A. Erturk, Y. Wang, M. Sheng, Local pruning of dendrites and spines by caspase-3-dependent and proteasome-limited mechanisms. J Neurosci 34, 1672–1688 (2014).

41. D. J. Simon et al., A caspase cascade regulating developmental axon degeneration. J Neurosci 32, 17540–17553 (2012).

42. Z. Li et al., Caspase-3 activation via mitochondria is required for long-term depression and AMPA receptor internalization. Cell 141, 859–871 (2010).

43. P. E. Mouser, E. Head, K. H. Ha, T. T. Rohn, Caspase-mediated cleavage of glial fibrillary acidic protein within degenerating astrocytes of the Alzheimer’s disease brain. AmJ Pathol 168, 936–946 (2006).

44. L. Acarin et al., Caspase-3 activation in astrocytes following postnatal excitotoxic damage correlates with cytoskeletal remodeling but not with cell death or proliferation. Glia 55, 954–965 (2007).

45. R. Aras, A. M. Barron, C. J. Pike, Caspase activation contributes to astrogliosis. Brain Res 1450, 102–115 (2012).

46. S. Villapol, L. Acarin, M. Faiz, B. Castellano, B. Gonzalez, Survivin and heat shock protein 25/27 colocalize with cleaved caspase-3 in surviving reactive astrocytes following excitotoxicity to the immature brain. Neuroscience 153, 108–119 (2008).

47. B. Hu et al., The DNA-sensing AIM2 inflammasome controls radiation-induced cell death and tissue injury. Science 354, 765–768 (2016).

48. A. Di Micco et al., AIM2 inflammasome is activated by pharmacological disruption of nuclear envelope integrity. Proc Natl Acad Sci U S A 113, E4671–4680 (2016).

49. A. Dumas et al., The inflammasome pyrin contributes to pertussis toxin-induced IL-1beta synthesis, neutrophil intravascular crawling and autoimmune encephalomyelitis. PLoS Pathog 10, e1004150 (2014).

50. E. G. O’Koren, R. Mathew, D. R. Saban, Fate mapping reveals that microglia and recruited monocyte-derived macrophages are definitively distinguishable by phenotype in the retina. Sci Rep 6, 20636 (2016).

51. L. M. Holt, S. T. Stoyanof, M. L. Olsen, Magnetic Cell Sorting for In Vivo and In Vitro Astrocyte, Neuron, and Microglia Analysis. Curr Protoc Neurosci 88, e71 (2019).

52. K. J. Livak, T. D. Schmittgen, Analysis of relative gene expression data using real-time quantitative PCR and the 2(-Delta Delta C(T)) Method. eMthods 25, 402–408 (2001).

